# Alternative Import-Channels And Destinations Of Mitochondrial PINK1 Controlled By Trans-Membrane-Domain Structural Plasticity

**DOI:** 10.1101/2024.11.06.622366

**Authors:** James S. Lorriman, Adam G. Grieve, Robin A. Corey, Ian Collinson

**Affiliations:** School of Biochemistry, University of Bristol, Bristol, BS8 1TD, UK; School of Physiology, Pharmacology and Neuroscience, University of Bristol, BS8 1TD, UK

## Abstract

Entry of the PINK1-kinase into the mitochondrial inner-membrane results in cleavage by the rhomboid-protease PARL, followed by retro-translocation back to the outer-membrane and proteasomal-degradation. Failure of this process, in compromised mitochondria, leads to kinase activation at the surface, and ultimately mitophagy. Analysis of PINK1-import within intact cells reveals an alternative pathway into the matrix. Structural modelling predicts that PINK1’s trans-membrane-domain (TMD) forms either an α-helix or α/β-hybrid at the interface between Tim17, of the TIM23-core-complex, and, respectively, either Romo1 or PARL. These mutually exclusive interactions both encapsulate a hydrated protein-channel. The α-helical-TMD form adopts a pose suggestive of translocation through the Romo1/Tim17-channel, while the α/β-hybrid-TMD is retracted into PARL’s active-site for cleavage, presumably after import through the adjacent rhomboid/Tim17-channel. We propose structural plasticity of the PINK1-TMD underlies alternative destinies for full-length matrix-import or cleavage/retro-translocation. The results reveal new insights of PINK1’s role in mitochondrial function, quality-control and early-onset Parkinson’s disease.

## Introduction

Nearly all mitochondrial proteins are synthesised in the cytosol, save only a few (13 in humans) which are encoded by the mitochondrial genome. Consequently, there is a high demand for protein import in mitochondrial biogenesis, maintenance and regeneration. Proteins destined for import contain mitochondrial targeting sequences (MTS) which direct them to the mitochondrial outer-membrane, inter-membrane space (IMS), inner-membrane or matrix ^1,2^. Those heading for the matrix are usually (but not always) delivered to the mitochondria as precursor proteins with a cleavable N-terminal MTS (pre-sequence) ^3^.

Import begins with targeting to the translocase of the outer-membrane (TOM)–acting as both receptor and protein-channel for entry into the IMS ^4^. Translocases of the inner-membrane (TIM) are then required for transport across and into the inner-membrane ^5,6^. The major route into the matrix occurs *via* the TIM23-pathway, in a process characterised by two distinct rate-limiting steps: (1) transport through TOM, and (2) initiation of transport through the TIM23-complex ^7^. Subsequent translocation into the matrix then proceeds very quickly (non-rate limiting). Passage across and into the inner-membrane are both driven by the membrane potential (Δψ), while matrix entry also requires ATP turnover by the mitochondrial Hsp70 homologue (mtHsp70).

The core TIM23-complex comprises the homologous subunits Tim23 and Tim17, of which humans have two orthologs (TIMM17A and TIMM17B), along with Tim44 peripherally associated on the matrix side. The latter associates with mtHsp70 for ATP-driven translocation ^8–11^. It had been assumed that the protein-channel is formed at the interface between Tim17 and Tim23, however when the structure of the core-complex was resolved, it did not contain a hydrophilic water containing pore ^12^. Instead, Tim17 forms a hydrophilic ‘slide’, similar to that found in the YidC-family ^13,14^; perhaps suitable for the insertion of proteins into the membrane (like YidC) ^15^, but not for conducting polypeptides across the membrane.

Formation of a protein-channel is likely to require association with additional proteins; for example Mgr2 in yeast (Romo1 in humans) ^12,16,17^; AlphaFold modelling suggests that this is indeed the case ^12,16^. Though, one problem with this proposal is that neither of Mgr2 and Romo are essential ^18,19^. Once the channel is formed then translocation necessarily occurs with minimal leakage of small molecules, to conserve the proton motive force (PMF) across the inner-membrane. This is presumably accomplished through an intimate association of channel constituents and the precursor, and by fast passage of precursors across the inner-membrane, as noted ^7^. Membrane integrity may also be achieved by transient assembly of the protein-channel upon demand.

One mitochondrial-targeted protein requiring this machinery is the stress-responsive protein PTEN-induced kinase 1 (PINK1). PINK1 is a mitochondrial serine-threonine kinase most widely known for its role in targeting damaged mitochondria for degradation *via* its interaction with the ubiquitin ligase Parkin ^20–22^. This process, termed mitophagy, ensures the maintenance of a functional cell-wide mitochondrial network, with errors in the process being linked to Parkinson’s Disease. PINK1 comprises an N-terminal MTS, an outer-membrane targeting sequence (OMS) followed by a transmembrane domain (TMD), and a C-terminal kinase domain (Figure 1a).

**Figure 1:**
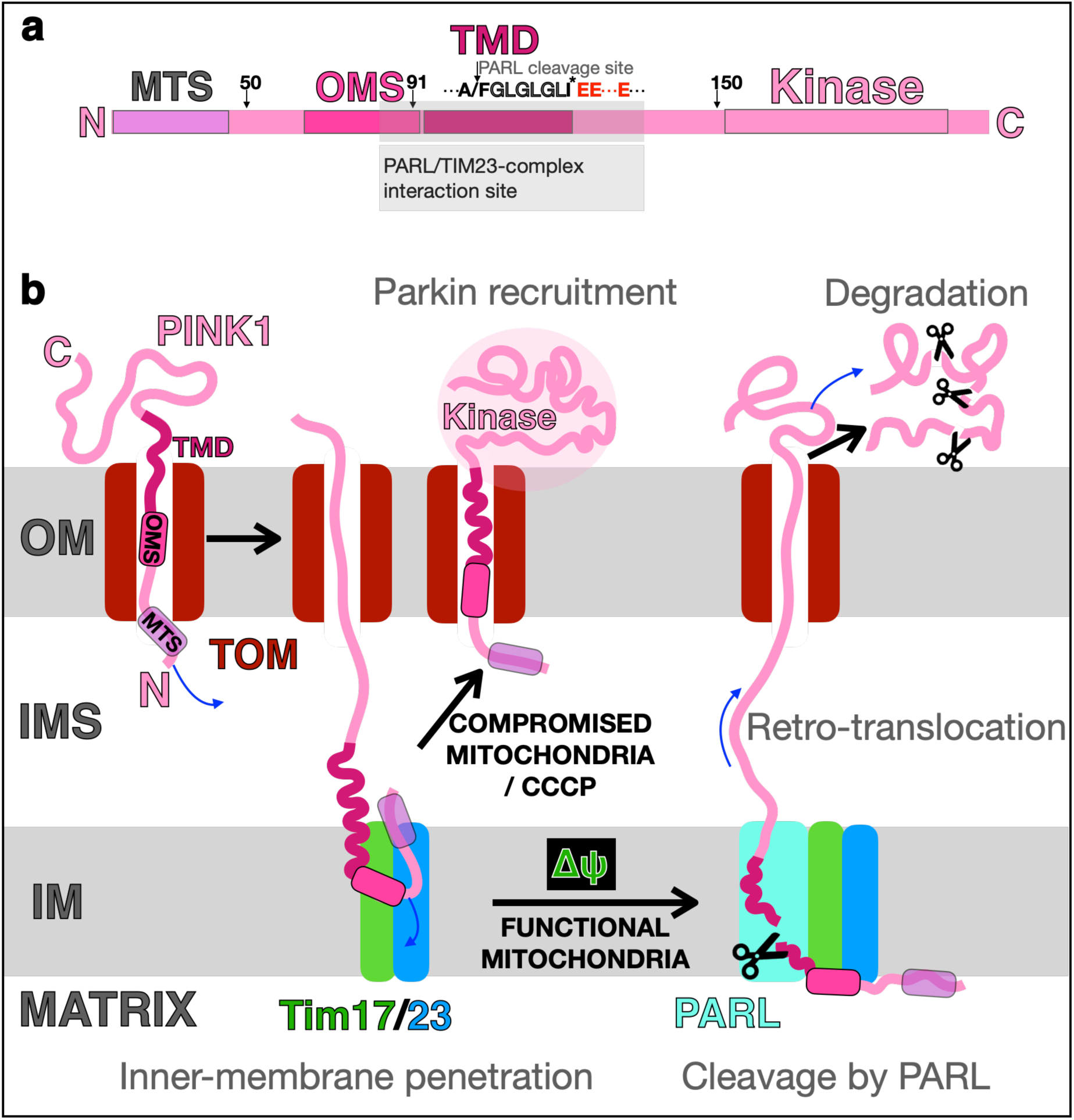
Domain structure and mitochondrial import of human PINK1 (human). **a.** Organisation of PINK1 showing: the MTS, mitochondrial targeting sequence; OMS, outer mitochondrial membrane localization signal; TMD, trans-membrane domain and the kinase domain. Key amino-acid sequence shown for the PARL cleavage site, GLGLGL-motif, I111 and conserved glutamates (see text). The N-terminal truncation sites at 50, 91 and 150 residues are shown (see Figure 4f-h) and the region interaction with PARL and the TIM23-complex are shaded grey (see Figure 4). **b.** Schematic representation of the PINK1 import system; see text for further details. Key elements of PINK1 are labelled (top left, as in **a**); mitochondrial targeting sequence (MTS), outer-membrane targeting sequence (OMS), trans-membrane domain (TMD), with consistent colouring throughout; blue arrows indicate the direction of protein translocation. With functional mitochondria, PINK1 is thought to undergo partial import into mitochondria followed by cleavage within the transmembrane domain by the inner-mitochondrial membrane resident rhomboid protease PARL. PARL cleavage precedes a process of retro-translocation, whereby PINK1 slides back out into the cytoplasm and is subject to degradation by the proteasome. In a compromised situation, PINK1 is stabilised at the outer membrane whereupon its dimerisation, mediated by the TOM-complex, facilitates autophosphorylation and the subsequent onset of mitophagy.

In normally functioning mitochondria, it is thought that PINK1 partially imports into mitochondria, with the N-terminal region (MTS, OMS and TMD) sitting deeply within the inner-membrane. This then enables cleavage between the OMD and the TMD (between residues A103 and F104) by the inner-membrane resident, the rhomboid protease PARL (Figure 1b) ^23–25^. The bisected TMD and kinase of PINK1 then undergo retro-translocation out of the mitochondria for degradation by the proteasome; whereby the new N-terminus (F104) acts as a type-2 N-degron, recruiting E3 enzymes UBR1, UBR2 and UBR4 that target the protein for degradation ^26,27^. The cleaved MTS, OMS and other half of the TMD remain in the inner-membrane; presumably this fragment is eventually released from the TIM23-PARL complex for degradation.

Conditions of mitochondrial stress, i.e. membrane depolarisation (induced also artificially by the ionophore CCCP), results in import arrest and, as a result, failure of PINK1 inner-membrane insertion and PARL cleavage (Figure 1b). In these circumstances, PARL is not recruited, and the uncleaved full-length PINK1 protein defaults to the outer-membrane; so the kinase domain is stabilised at the mitochondrial surface ^28^. Subsequent PINK1 dimerization facilitates autophosphorylation and Parkin recruitment ^29,30^. Phosphorylation of Parkin then induces a downstream ubiquitination cascade, leading to proteasomal-mediated protein degradation and ultimately mitophagy ^31,32^. It has also been reported that reactive oxygen species (ROS) also promote PINK1/ Parkin dependent mitophagy ^33^.

These alternative scenarios explain how PINK1 acts as a reporter of mitochondrial fitness for quality control. However, the current model does not consider the possibility and consequences of fully imported PINK1, where the kinase domain crosses the inner-membrane to enter the matrix ^34^. This prospect has been suggested by the identification of a potential PINK1-kinase substrate within the mitochondrial interior. PINK1 knock-outs result in a loss of phosphorylation of the matrix facing subunit NdufA10 of the respiratory Complex I, required for high rates of electron transfer ^35^. Additionally, it has been shown that PINK1 activity is regulated through degradation by the Lon-protease in the matrix ^36^. These observations suggest that, in addition to stress-related mitophagy signalling, PINK1 has additional regulatory roles relevant to mitochondrial structure, function and homeostasis. This could be achieved indirectly through trans-membrane signalling cascades from the cytosolic kinase domain of PINK1 to the IMS and matrix. Alternatively, regulation of Complex I, *inter alia*, could be activated directly by matrix localised PINK1.

Given uncertainties about these alternative destinations of PINK1 we re-examined PINK1 mitochondrial transport, including its potential for both IMS and matrix import. For this, we had to overcome considerable technical barriers. The first was the production of full-length human PINK1, and the second the deployment of an accurate and real-time cellular assay for mitochondrial import (MitoLuc) ^37^. The assay was further developed in this study to monitor transport into the IMS as well as the matrix. Thereby, we were able to construct and purify a series of human PINK1 precursor proteins amenable for the analysis of its import. The import data are supported by structural modelling (using AlphaFold2 ^38–40^) and molecular dynamics (MD) simulations that address the mechanistic basis for PINK1 inner-membrane insertion for proteolysis and transport.

The combined results reveal that PINK1 can be directed either into the matrix or the outer-membrane and suggest a mechanism for this discrimination, based around the structural plasticity of PINK1’s TMD. These data, along with the effects of Parkinson’s disease-causing residue substitutions on the import behaviour of PINK1, have motivated a re-evaluation of PINK1’s role in mitochondrial regulation. The results also raise the prospect of there being multiple bespoke protein-channels through the inner-mmebrane for different purposes, formed by the assembly of the TIM23-core-complex with various accessory partners, with Mgr2/Romo1 being one of many alternatives.

## Results

### Import of PINK1 into the IMS occurs in 2 rate limiting steps

To better understand the import behaviour of PINK1, we first set out to interrogate its transport across the outer-membrane into the IMS. This was achieved by the development of our in-cell import assay–MitoLuc ^37^. This involved the encapsulation of a large fragment of a split luciferase (11S) into the mitochondrial IMS (i11S) of intact human cultured cells; while the corresponding small fragment (pep86) is incorporated into the import protein (PINK1). The cells (in this case HEKs) are then perforated to allow the entry of the import protein substrate and assay reagents. Successful import then results in the association of the small and large fragments for the formation of an active luciferase for a real time and accurate monitor of transport (Figure 2a; i11S); see Materials and Methods for details.

**Figure 2:**
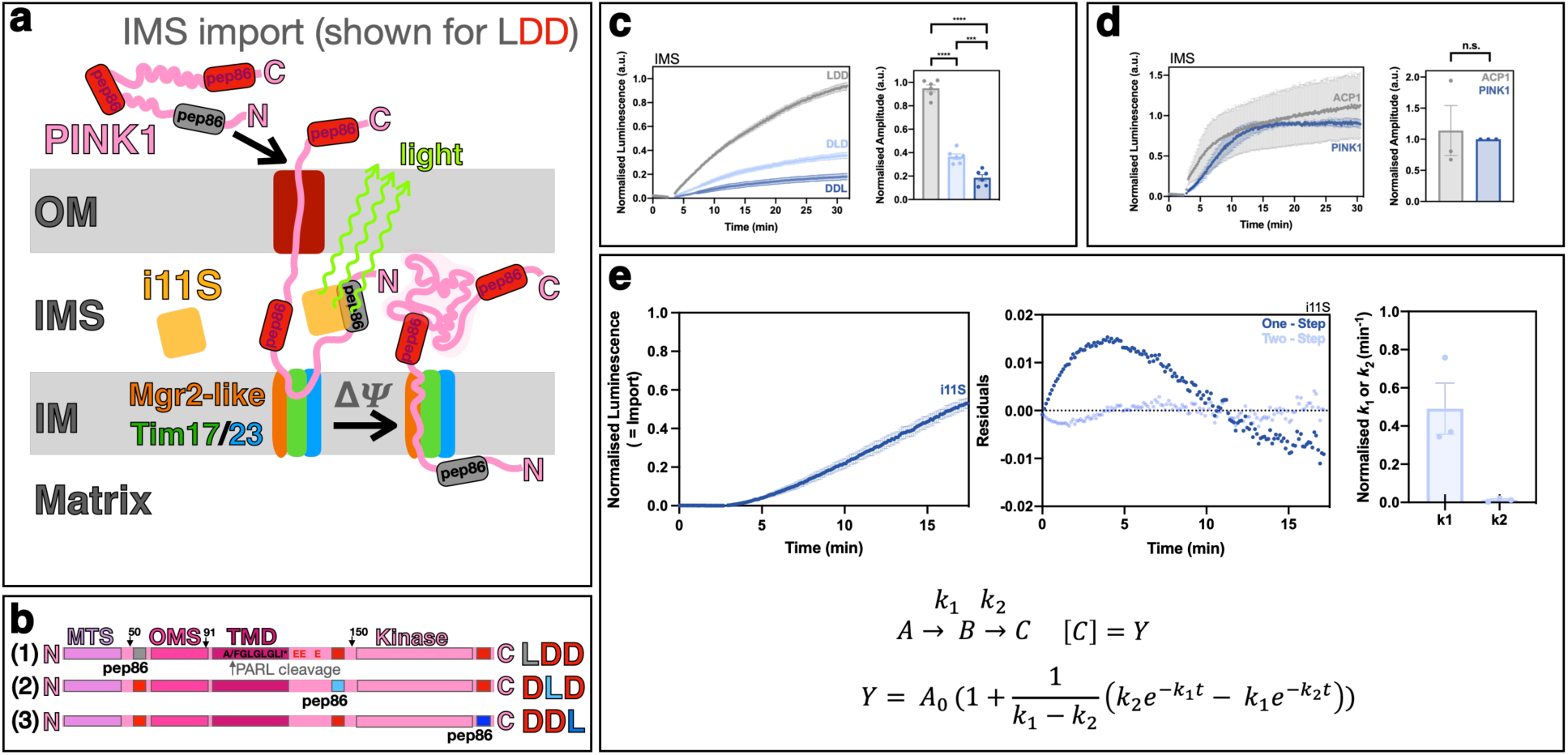
Import of PINK1 into the mitochondrial IMS proceeds in two distinct rate-limiting steps. **a.** Schematic representation of the MitoLuc import assay–monitoring entry into the IMS by pep86 association with IMS targetted 11S (i11S) for the formation of an active luciferase. PINK1 is depicted with 3 pep86 luciferase fragments. Separate experiments were conducted with the PINK1 LD series of 2 inactive (‘dark’, D) and 1 active (‘light’, L) pep86, in sequence: **L**DD, D**L**D and DD**L**; the former being shown in this schematic (see text for further details). **b.** The PINK1 LD series (**L**DD, D**L**D and DD**L**) generated for assessing mitochondrial IMS and matrix (Figure 3) import of PINK1– labeling as is Figure 1a. **c.** MitoLuc traces for import of the PINK1 LD series into the IMS, in each case background luminescence was recorded for prior to addition of each of the PINK1 precursors. Normalised amplitudes were calculated from the point at which traces plateaued and plotted for each LD precursor. Error bars represent SEM and data represent an N=3-5 biological repeats. A paired t-test was used to determine significance of the normalised amplitudes. **d.** MitoLuc trace and associated normalised amplitudes for import of PINK1 LDD and the matrix-targeted precursor ACP1 into the IMS. **e.** The MitoLuc data for the PINK LDD variant (upper left) was fitted to one-step (equation not show) and two-step (lower) models for import into the IMS. Residual plots (upper middle) represent the difference between the experimental data and fit. Normalised rate constants k_1_ and k_2_ values (upper right) obtained from fitting the IMS import data to the two-step model. Data shown represent N=3, error bars show SEM.

To provide a more granular readout of the import process, we engineered the luminescent reporter (pep86) in 3 different positions along PINK1: (1) at the N-terminal domain between the MTS and OMS, (2) between the TMD and the kinase domain and (3) at the C-terminus (Figure 2a, 2b and S1). The 3 variants all contained one active luminescent ‘light’ (L) and two inactive ‘dark’ (D) pep86 sequences; the latter disabled by pairwise amino acid swapping (x3). Thus, the three construct, hereafter termed LDD, DLD and DDL, were identical save for the position of the active reporter.

The full-length proteins were successfully purified (typical example shown in Figure S2) and shown to associate with the large fragment of the luciferase (11S) in solution for the formation of an active luciferase and the liberation of bioluminescence (Figure S3). Based on our previous analysis and methodological development ^7,41^, we know that these association kinetics are suitable for implementation in the MitoLuc assay. This is because the internalised mitochondrial volumes are tiny, so the concentration of pep86 and 11S will be very high (much higher than those shown in Figure S3). Therefore, the association rates will be very high (substancially higher than the rate of import) and non-rate limiting. Therefore, in the MitoLuc assay, the luminescence response reports on the rate of mitochondrial import, rather than on the assembly of 11S and pep86. Slight variations in the bioluminescent yield of the various PINK1-pep86-11S complexes (Figure S3, B_max_) were used to scale the subsequent import measurements accordingly.

Progression of the 3 reporting regions of PINK1 into the IMS within HEK cells was then monitored over time, one by one, for each of the LDD, DLD and DDL variants. Figure 2a describes the import assay for LDD, whereby entry of the N-terminal region into the IMS is monitored. The DLD variant reports on the entry of the region prior to the kinase domain into the IMS, and DDL the C-terminus as a result of IMS entry of the full-length protein including the kinase domain. The luminescence traces (Figure 2c) chart the progression of the respective regions across the outer-membrane. The amplitude (maximum luminescence) reflects the total amount of the pep86 reporter entering the IMS and the rate reflects the time taken to cross the outer-membrane. Both of these were similar in PINK1 to the canonical matrix-bound precursor protein (ACP1) characterised previously ^37^ (Figure 2d).

Import of ACP gives us a measure of IMS build up of a precursor on its way to the matrix. This is as expected because we have previously shown that initiation of transport across the inner-membrane is rate limiting ^7^. In the case of PINK1, from this data alone, we cannot say if (or how much) of the IMS signal is due to its final destination in the IMS, or as a result of build up of proteins on their way to the matrix.

The PINK1 data could be fitted to a two-step reaction mechanism (Figure 2e) for transport across the outer-membrane. The rates are unaffected by the position of the luciferase fragment within PINK1 (Figure S4), demonstrating that the passage of the N-terminal, central and C-terminal regions of PINK1 into the IMS occurs by the same mechanism. Presumably, these two steps are: (1) association with the TOM complex and (2) passage through the channel into the IMS.

The amount of material in the IMS decreases progressively from the N-terminal to the central and C-terminal regions (Figure 2c), which probably reflects the directionality of transport (N→C-terminus) through TOM40, as well as the relative slow rate (rate-limiting) of this step ^7^. Transport quantities into the IMS were insensitive to respiratory and ATP synthase (inner-membrane) inhibitors (Antimycin A and Oligomycin, ‘AO’) (Figure S5); this was as expected because the outer-membrane is not subject to their effects.

### PINK1 is fully imported into the matrix

Following import across the outer-membrane and into the IMS it is generally accepted that PINK1 becomes partially incorporated into the inner-membrane of uncompromised mitochondria *via* the TIM23-complex ^25^; whereafter it associates with PARL (Figure 1b). It has been suggested, though not widely accepted, that the interaction with the import machinery can also promote PINK1 entry into the matrix ^35,36,42^.

To explore the prospect of partial or complete transport of PINK1 across the mitochondrial inner-membrane, we interrogated our PINK1 LDD, DLD, and DDL variants using the MitoLuc assay monitoring precursor entry into the matrix (Figure 3a) with 11S encapsulated in the matrix (m11S). The data reveal that PINK1 does indeed enter the mitochondrial matrix (Figure 3b). Furthermore, the luminescence signal from the C-terminal pep86 reporter demonstrates that PINK1 is fully imported, *i.e.* the entire protein crosses the inner-membrane to enter the matrix (Figure 3b; DDL).

**Figure 3:**
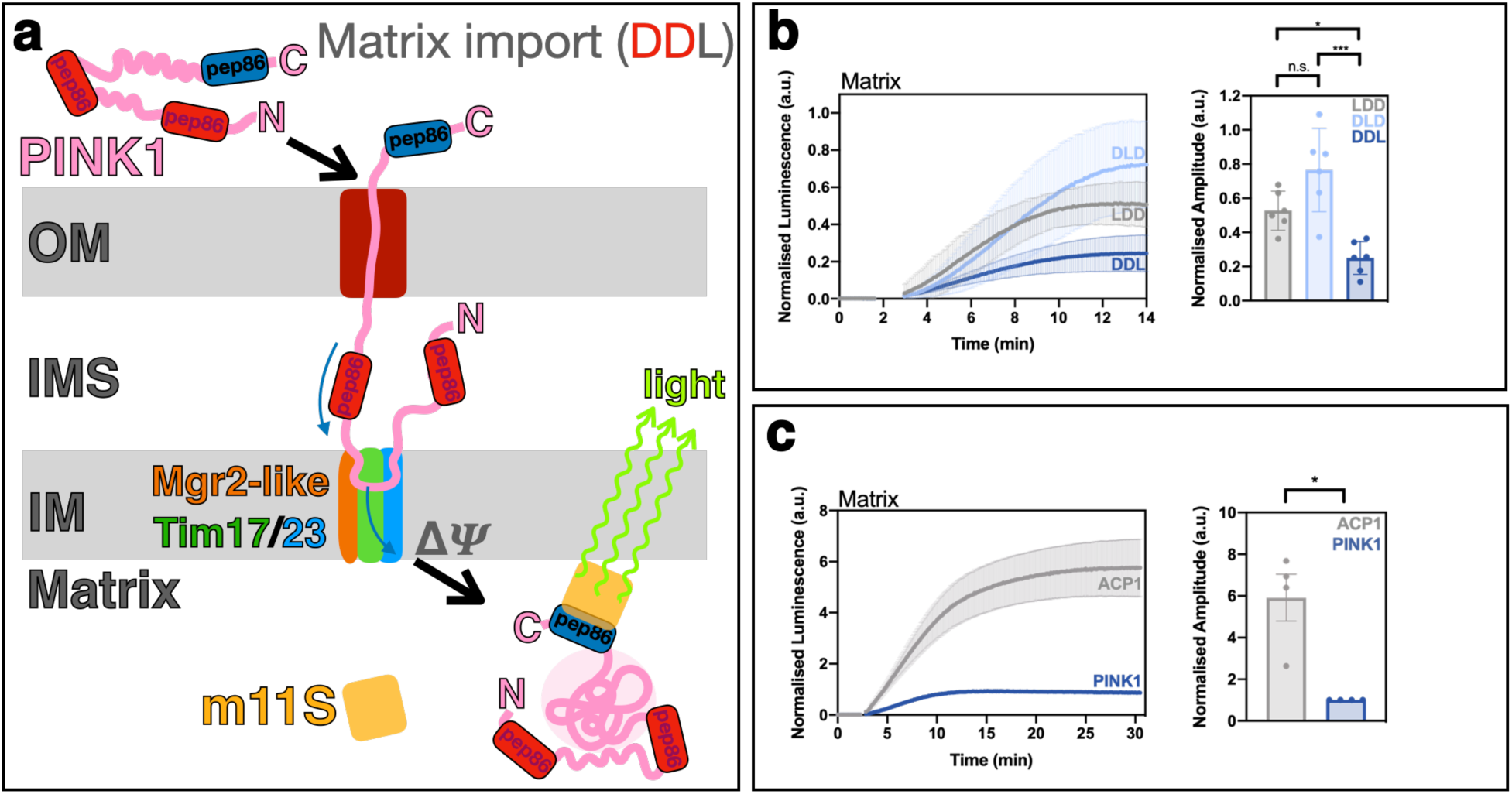
PINK1 is fully imported into the mitochondrial matrix. **a.** Schematic shown as in Figure 2 for monitoring import into the matrix, rather than the IMS. In this depiction the import of the PINK1 variant DD**L** (Figure 2) is shown. **b.** MitoLuc trace for the import of the PINK1 LD series (Figure 2b; **L**DD, D**L**D and DD**L**) into the matrix, in each case background luminescence was recorded prior to addition of the specific PINK1 precursor variant. Normalised amplitudes were calculated from the point at which traces plateaued and plotted for each of the LD precursors. **c.** MitoLuc traces and associated normalised amplitudes for the import of PINK1 DDL and the precursor ACP1 into the matrix.

Unlike the IMS data, which fitted to a two step model, the matrix import datasets are too complex to fit to a kinetic model. Nevertheless, the data reveals several basic features about PINK1’s mitochondrial trafficking. While substantial, the total quantity of import is somewhat lower than the canonical matrix-targeting benchmark ACP1 (also carrying the luminescence reporter pep86 at the C-terminus; Figure 3c), suggesting that not all of the PINK1 enters the matrix, and that some of it is retained in the IMS. Further, in contrast to IMS import, subsequent passage into the matrix is sensitive to Antimycin A and Oligomycin (Figure S6), suggesting that, as expected, translocation occurs *via* the Tim23 pathway, which requires both ATP and the membrane potential (Δψ) ^7,10,11,43,44^. Critically, deletion of the N-terminal targeting region of PINK1, up to the kinase domain, completely ablates the luminescent signal (see below). This demonstrates that the assay is a genuine measure of matrix import, rather than a result of PINK1 association with residual 11S in the cytosol.

### Modelling of PINK1 and its interaction partners reveals structural plasticity in the PINK1 TMD

The results above demonstrate that PINK1 can be fully transported across the inner-membrane. Therefore, following entry into the IMS and contact with the inner-membrane PINK1 has alternative destinations in functional mitochondria: (1) delivery to the rhomboid protease PARL for cleavage and retro-translocation to the outer-membrane, as previously described) ^23–25^, or (2) transport into the matrix. We presumed both to be mediated through interactions with the TIM23-complex. To better understand these distinct pathways, and how they are discriminated, we explored the predicted structures of PINK1 in complex with PARL, both alone and in combination with the TIM23-core-complex.

Structural models of PINK1 were produced by the deep learning AlphaFold2 (AF) program, see Methods for full details. Details of all AF models produced for this study are given in Supplementary Table 1. Top ranking models and plots of quality metrics are available for download from https://osf.io/xj9ca/, with key plots also included in the supplementary figures.

Initially, an AF model of the PINK1-PARL complex was built, with the highest scoring pose shown in Figure 4a. Each chain of the model is in good agreement with experimentally determined structures: the kinase domain of the PINK1 model aligns well with the resolvable region of *Pediculus humanus corporis* PINK1 (residues 147-574) ^45^, with an RMSD of 0.11 nm (Figure S7a). The PARL chain of the model compares well with the *E. coli* rhomboid homologue GlpG ^46^ (Figure S7a), with slightly higher structural variability than the PINK1 chain (RMSD 0.64 nm), expected due to the vast evolutionary distance between human PINK and *E. coli* GlpG. Accordingly, the predicted Local Distance Difference Test (pLDDT) is generally favourable, at ca. 80-90 for most of each chain Figure S7b. The pTM score across the complex is respectable at 0.63.

**Figure 4:**
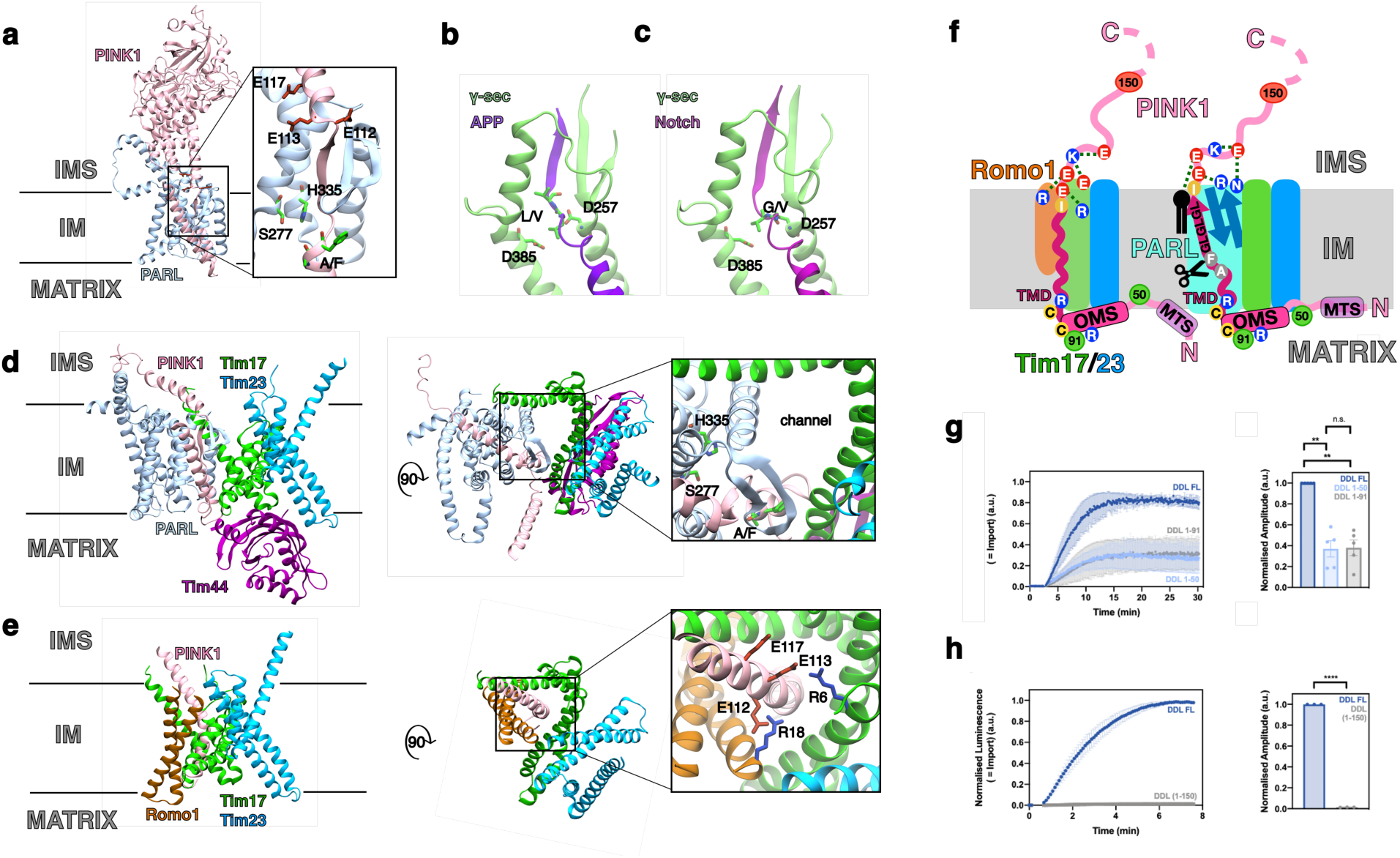
AlphaFold2 modelling of PINK1 and its interactions with PARL and the TIM23-core-complex. **a.** Top ranked of five AF models of the PINK1-PARL complex. Inset: Close up of the PINK1-PARL β-sheet (BS), showing the proximity to the PINK1 cleavage site between A103 and F104 (A/F) and conserved three glutamates (E112, E113 and E117) residues, as well as the active site catalytic residues of PARL (S277 and H335). **b.** and **c.** Respective views of γ-secretase bound to amyloid precursor protein (APP; PDB 6IYC) and Notch (PDB 6IDF). The active site aspartate residues are shown (D257 and D385), as well as the cleavage site, between leucine and valine (L/V) for APP and glycine and leucine (G/V) for Notch. **d.** View of the hybrid AF model of the PINK1-PARL-Tim17-Tim23-Tim44 complex, see methods for modelling details. PINK1 has been truncated after residue 150 for clarity. The complex is shown side on from the membrane (left) and from the IMS (right). Inset: zoom of the potential protein-conducting pore, formed between Tim17 and PARL, the adjacent PARL active site (S277 and H335) and the cleavage site of PINK1 (A/F). The lines indicate the position of the bilayer. **e.** Same views as in panel D of the AF model of Tim17-Tim23-Romo1 and the PINK1 TMD. Inset: a zoom on the PINK1 TMD, with the functionally important acidic residues highlighted (E112, E113 and E117). The lines indicate the position of the bilayer. **f.** Schematic of the N-terminal regions of PINK1 associated with the TIM23-core-complex including either Romo1 (left) or PARL (right). Key features include the MTS, OMS and TMD of PINK1, highlighting: the GLGLGL-motif, the three conserved glutamates (E), the PARL cleavage site (A/F), the N-terminal truncation sites 50, 91 and 150 (see below g and h), conserved cysteines and arginines (2Cs and 2Rs) described later (Figure 5) and I111 (see later Figure 7). In the PARL associated complex the rhomboid’s two β-strands are shown augmented by a third (GLGLGL of PINK1). Key interactions of the 3Es are shown with Romo1 and Tim17 (left): including arginines (R) of Romo1 and Tim17, as well as a glutamate (E) of Tim17, *via* an intra-molecular contact with a PINK1 lysine (K)–(see also **e,** and later Figure 6e). In the complex associated with PARL (right) the 3Es are shown interacting with arginine (R) and asparagine (N) residues of PARL, as well an intra-molecular interaction lysine (K), and with a the head group of a phospholipid–(see also later Figure 5e). **g.** MitoLuc trace for the import of the PINK1Δ(1-50) and PINK1Δ(1-91) truncated precursors into the matrix compared to the full-length (FL) protein. Normalised amplitudes were calculated at the point of plateau and plotted. **h.** MitoLuc trace for import of PINK1Δ(1-150) into the matrix relative to the FL precursor. Normalised amplitudes are plotted.

In general, the predicted aligned error (PAE) score between the PINK1 and PARL chains in the complex ‘is low. This is expected, as any complex formed by these proteins is likely be transient. Similarly, whilst not so low as to reject the model, the ipTM is moderately low at 0.5. However, a notable high scoring PAE region can be seen in the graph (Figure S7c, arrow), indicating a more confident region of the model. This region corresponds to a small three stranded β-sheet formed between PINK1 and PARL (Figure 4a). One strand of the β-sheet is formed from a section of the TMD region of PINK1 (sequence GLGLGL) between the cleavage site and three highly conserved acidic residues (Figure 4a), while the other two strands are contributed by PARL.

Thus, in the presence of PARL, the PINK1 TMD spans the membrane as an α/β-hybrid. Interestingly, in the AF model of PINK1 alone taken from the EBI database ^47^, the TMD including the GLGLGL-motif is entirely α-helical (Figure S7d). Notably, the α/β-hybrid arrangement brings the PINK1 cleavage site into proximity of PARL protease active site (Figure 4a: S277/H335). This configuration shows strong similarities to the structure of γ-secretase bound to amyloid precursor protein (APP) and to Notch ^48^, including the proximity of the active site to the substrate, and the formation of the three-strand β-sheet (Figure 4b-c). This supports the designation of this model as a plausible pre-cleavage PINK1/PARL interaction. In the modelled complex the kinase domain is shown in the IMS, as predicted by AF (Figure 4a), though it is unclear if PINK1 ever folds in the IMS, or if it remains unfolded prior to transport to the matrix or retro-transport the outer-membrane.

### Structural modelling identifies potential alternative protein-channels at the interface between the TIM23-core-complex and either Romo1 or PARL

To investigate the structure of PINK1 in the context of the mitochondrial import machinery, we augmented the AF PINK1-PARL model to also include the TIM23-core-complex (Tim17-23-44) (Figure 4d). This was achieved by creating an AF-model of the latter with PARL bound (Tim17-23-44-PARL). This model had solid AF scores of pTM=0.66 and ipTM=0.65. The pLDDT was high for the whole complex (Figure S8a), except for the N-terminal region of Tim44 which we excluded from downstream analyses. There was a good PAE score between PARL with Tim17, Tim23, and the C-terminal region of Tim44 (Figure S8a), suggesting a confident model prediction. The Tim17-23-44 arrangement closely resemble the published cryo-EM structure of the yeast complex (Figure S8b ^12^), with an RMSD of 0.15 nm between them. The PINK1-PARL and Tim17-23-44-PARL models were combined by overlaying the PARL subunits of the two complexes to create a predicted structure of Tim17-23-44-PARL-PINK1.

The resulting model reveals formation of a putative protein channel between Tim17 and PARL situated immediately adjacent to the rhomboid’s proteolytic active site (Figure 4d: S277 and H335). This channel closely resembles the proposed protein pathway formed between Tim17 and Mgr2 ^12^ (Figure S8d). Here, the TMD of PINK1 is incorporated into the PARL active site, adjacent to–but not within–the described Tim17 protein-channel. This channel is also evident in the model of the Tim17-23-44-PARL complex, produced without PINK1 (Figure S8c).

In the yeast Tim17-23-44 cryo-EM study, the authors used AlphaFold2 to predict a protein channel between Tim17 and Mgr2 ^12^, which we were able to reconstruct in AlphaFold2 (Figure S8d; pTM=0.69, ipTM=0.74). For the human conterparts, it was also possible to model Tim17-23-44 complex with Romo1, the homologue of Mgr2 ^49,50^, (Figure S8d; pTM=0.79, ipTM=0.76). This model is in agreement with a recent study, which also showed that the Romo1 interaction can be formed by both orthologs of Tim17 (TIMM17A and TIMM17B) ^16^.

The position of Mgr2 and Romo1 overlays with our predicted PARL binding site within the complex (Figure S8c, S8d), suggesting there is a mutual exclusivity between Mrg2/Romo1 and PARL. Based on this good agreement between Romo1 and PARL placement in the TIM23-complex, we modelled the N-terminal domain of PINK1 into the Tim17-23-Romo1 complex (Figure 4e). Here, the PINK1 N-terminal region is incorporated into the protein-channel at the interface between Tim17 and Romo1, positioned directly next to three acidic residues, known to be important for protein import (Figure 4e) ^17^. Interestingly, the PINK1 TMD is modelled as an α-helix rather than the α/β-hybrid of the PARL complex. Therefore, it seems that Romo1 promotes α-helical TMD formation and transport of PINK1 into the matrix by the classical TIM23 translocation pathway, while PARL favours the α/β-hybrid conformation as a potential precursor to proteolysis (Figure 4f).

### Mitochondrial matrix import requires the TMD of PINK1

The AF models highlight the importance of the N-terminal region of PINK1 for inner-membrane association with both PARL and the TIM23-complex. The latter interaction predicts this region to be critical for matrix import. To examine this further, we constructed a series of successive N-terminal truncations of the PINK1 DDL variant for use in the MitoLuc assay: Δ1-50 (removal of the classical MTS), Δ1-91 (removal of the MTS and OMS) and Δ1-150 (removal of the MTS, OMS and TMD, retaining only the kinase domain) (Figure 1a, S1 and 4f). Again, minor variations in the modified proteins interaction with 11S and their luminescent yield were monitored (Figure S9) and used to scale the import analysis accordingly. The results show that import into the matrix is impaired (by ∼60%) in the Δ1-50 and Δ1-91 constructs (Figure 4g), but only fully abolished in the Δ1-150 construct (Figure 4h). Therefore, only the region between 91-150, including the TMD (containing PARL cleavage site), but not the OMS, is necessary and sufficient for association with the TIM23-complex and import. This is in excellent agreement with the AF models, which identify this region as being most intimately associated with the Tim17-23-44 model complexes, bound by either Romo1 or PARL.

### Molecular dynamics simulations identify interactions of PINK1’s TMD and a hydrated protein-channel potentially critical for inner-membrane translocation and cleavage

To further validate the AF modelling and explore the dynamics and interactions required for PINK1 cleavage, we carried out all atom molecular dynamics (MD) simulations of the PINK1-Tim17-23-44-PARL complex within a model mitochondrial membrane. During the simulations, the complex was very stable, with an RMSD for the transmembrane complex (excluding the PINK1 kinase domain) that plateaus at 0.38±0.02 nm for across the 5 repeats (Figure S10a). Crucially, the PINK1-PARL β-sheet was completely stable throughout the simulation; quantification of the hydrogen bonding revealed the average incorporation of ∼6 backbone hydrogen bonds inter-connecting the 3 strands (Figure 5a). Adjacent to this, the catalytic site of PARL and cleavage site of PINK1 are well solvated, with water molecules freely able to access and leave the region (Figure 5b). This is achieved in part through a constriction in the membrane around PARL (Figure 5c). In this configuration the PARL catalytic site remains close to the PINK1 cleavage site (A103/F104), typically a distance of about 0.65 nm; although in one instance it withdrawns to about 1.5 nm (Figure 5d, S10b).

**Figure 5:**
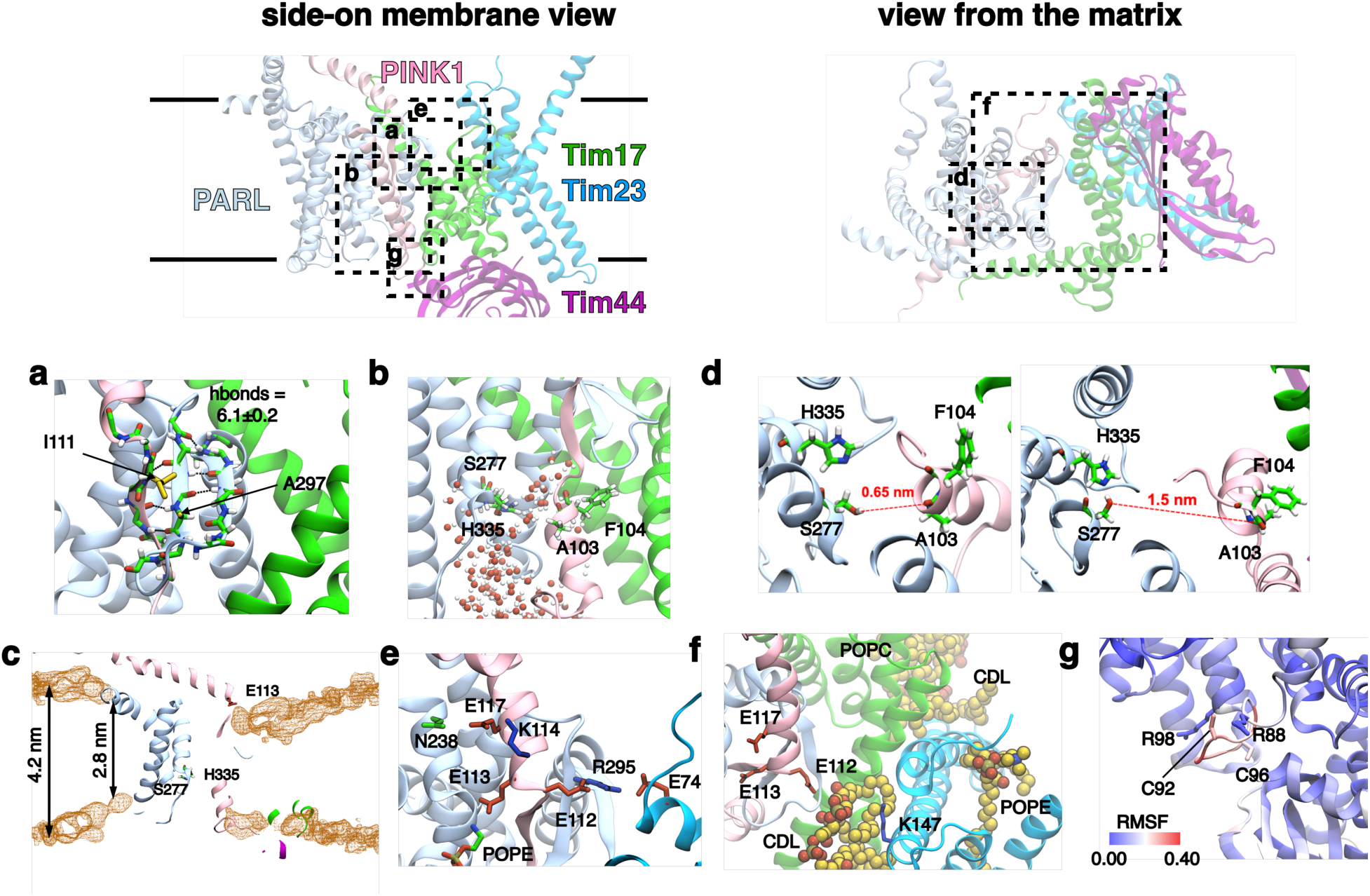
Molecular Dynamics analysis of the PINK1-PARL-Tim17-Tim23-Tim44 complex. **Upper panels:** views of the PINK1-PARL-Tim17-Tim23-Tim44 complex from the side (left; lines indicating the position of the bilayer) and view from the IMS (right) with PINK1 truncated after residue 150 for clarity. Boxes are drawn for orientation of panels **(a-g)**. **a.** View of the β-sheet formed by one β-strand of the PINK1 TMD and two from PARL post MD simulation. The backbone atoms of residues that contribute to the β-sheet are shown as sticks and coloured by atom type. Hydrogen bonds, as identified by VMD, are shown. The number of hydrogen bonds between the highlighted residues is denoted (mean and standard deviation), as computed using Gromacs over the final 200 ns of each simulation. The residues I111 and A297, which interact with one another, are shown as yellow sticks. **b.** View of the PARL active site, with catalytic residues (S277 and H335), and PINK1 cleavage site (A103/F104) post simulation, showing that the site is highly solvated. **c.** Showing the membrane thinning about the PARL protein. The density of all lipid phosphate atoms has been computed for a simulation fitted on PARL using the VMD VolMap tool and shown as orange mesh. For clarity, the density map has been clipped to only show a slice through the membrane around PARL. The PARL active site catalytic residues and E113 (of the three conserved Es) of PINK1 are shown. Approximate membrane thickness, taken from phosphate-phosphate distances are shown. **d.** Snapshot of the PARL active site and PINK1 cleavage site showing a typical (left, 0.65 nm) or maximum (right, 1.5 nm) distance from the hydroxyl of the catalytic residue S277 to the A/F cleavage site backbone. **e.** Highlighting the 3 conserved glutamates (E112, E113 and E117) of PINK1 and the interactions it makes with PARL (R295), Tim23 (E74), and a surrounding phospholipid (POPE). **f.** IMS view of the complex showing the most tightly bound lipids. Those on the righthand side were previously seen in the yeast Tim17/Tim23 structure. The CDL at the bottom is unique to the PARL-containing complex, and bridges PINK1, PARL, and Tim23. The interaction of K147 with CDL is of particularly high affinity, being occupied in 94% of all MD frames (as assessed using the PyLipID package ^66^). **g.** Root-mean-squared fluctuation (RMSF) analysis reveals the matrix loop containing Cys92 and Cys95 to be highly dynamic (units = nm).

Immediately following PINK1’s GLGLGL-motif of the PINK1/PARL β-sheet, there are 3 conserved glutamates known to be essential for normal PINK1 behaviour (Figure 2b, S1 and 4f). In our model and simulations, one of them, E112, interacts with R295 of PARL, which in turn interacts with E74 of Tim23 (Figure 5e), helping to stabilise the complex. The next one, E113, consistently binds to a POPE lipid, forming a crucial part of a high affinity lipid binding site (occupied by POPE for the entire simulation), which contributes to the membrane constriction around PARL (Figure 5c). Additional lipid binding sites were also noted in this vicinity, including cardiolipin (CDL), POPC and POPE, similar to those visualised in the yeast cryo-EM structure, as well as a novel CDL binding site between K147 of Tim23 and E112 of the PINK1/PARL β-sheet (Figure 5f). This suggests the CDL might be important for stabilising the PINK1/PARL/TIM-complex. The third glutamate, E117, forms interactions with K114 of PINK1 and N238 of PARL (Figure 5e), which would likely also contribute to inter-subunit stability.

Of further interest in this region is I111 of PINK1, found between the GLGLGL-motif and the 3 key glutamates. This forms a stable hydrophobic interaction with A297 of PARL in the middle strand of the β-sheet continuously throughout the simulation (Figure 5a). This interaction would be likely to stabilise the PINK1/PARL β-sheet. Of note, substitution of this residue to serine (I111S) is implicated in early onset Parkinson’s disease ^25^, suggesting a direct link of this region to a disease phenotype.

The simulated structure identifies an interesting consequence of the orientation of PINK1 within the PARL-Tim17-23-44 complex. A short loop between the TMD and OMS contains two conserved cysteines (C92 and C96), which poke into the matrix (Figure 4f, S1 and 5g). Both are near arginine residues (also conserved), which promote the conversion of thiols (–SH) to thiolates (–S^−^), and thereby enhance their sensitivity to oxidation ^51^. Therefore, they might be there for ROS sensing to monitor the activity of the electron transfer chain. In our simulations, these cysteines are very dynamic, which would fit this role (Figure 5g, S10c).

Critically, the simulations show that the interface between Tim17 and PARL is predicted to be full of water (Figure S11), and therefore ideally suited for protein translocation. For instance, for the entry of PINK1 into the membrane for proteolysis, as well as for the retro-translocation of the cleavage product.

### Molecular dynamics simulations identify a hydrated protein-channel at the Tim17/Romo1 interface stably occupied by the TMD of PINK1 for matrix-entry

We next assessed the nature of the PINK1 TMD within the Romo1-bound import complex. To allow longer simulations, we modelled only the TMD of PINK1 into the Tim17-23-Romo1 complex with Tim44 removed (Supplementary Table 1). To verify that a short region of PINK1 should be stable, we also truncated PINK1 in the cleavage intermediate complex (as above with the Tim17-23-44-PARL complex and with Tim44 removed) and ran simulations in parallel (Figure 6a, b). The data show that the truncated PINK1 TMD is stable within each of complexes; despite higher dynamics seen at the PINK1 termini– to be expected due to their artificially trimming–the central region is very stable.

**Figure 6:**
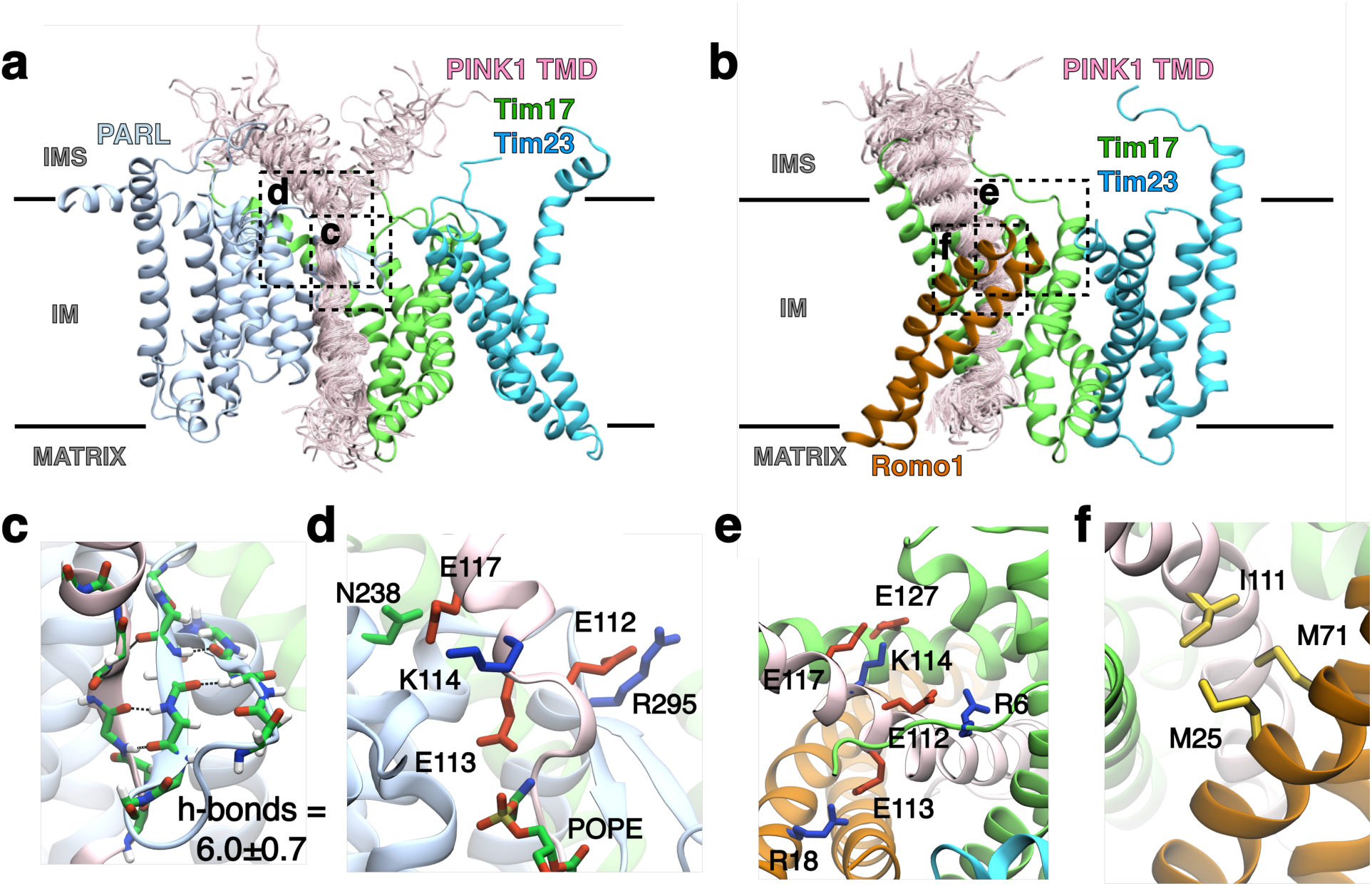
MD simulations of the TMD of PINK1 in the PARL and Romo1 containing TIM23-core-complexes. **Upper panels:** views of the MD simulations of the PARL-Tim17-Tim23 **(a)** and Romo1-Tim17-Tim23 **(b)** complexes, with the PINK1 TMD respectively in the proteolytic cleavage site (a), or the protein-channel (b), as positioned by AlphaFold2. The lines indicate the position of the bilayer. Multiple frames of the PINK1 TMD are overlaid to reveal the restricted dynamics of this polypeptide across the simulations. Orientation boxes are drawn for panels **(c-e)**. **c.** view of the β-sheet formed by the PINK1 TMD and PARL post MD simulation. As in Figure 5a (simulated with full-length PINK1), the backbone of the β-sheet is shown as sticks with hydrogen bonds shown as dashed lines and the computed average hydrogen bond number shown. **d.** Key interactions between the 3 conserved glutamates of PINK1 and PARL are preserved in the truncated TMD simulations (compared to Figure 5e). **e.** View of the Romo1-Tim17-Tim23-complex highlighting the interactiosn of the 3 conserved glutamates (see also Figure 4f). **f.** Highlighted interactions between I111 of PINK1 and Romo1 preserved during the MD simulations.

In the PARL-containing complex, the PINK1/PARL β-sheet remains as stable as in the simulations with full-length PINK1 (Figure 5a), again with ∼6 hydrogen bonds consistently formed between the three strands (Figure 6c). The RMSD for this central region of PINK1 in relation to the whole complex is 0.37±0.09 nm. PINK1 is even more stable in the Romo1-containing complex, with a RMSD of 0.20±0.02 nm. Moreover, the simulations highlight the very plausible existence of alternative protein-conducting channels forming between the TIM23-core-complex, specifically Tim17, and either PARL or Romo1 (Figure 6a, 6b and S11).

The simulations of the respective complexes bound to PARL or Romo1 presents compelling alternative structural views concerning the fate of PINK1: respectively, intermediates prior to either cleavage or matrix import. In the former, precisely consistent with the simulation containing full-length PINK1 (Figure 5e), the 3 conserved glutamates form stabilising interactions at the PINK1-PARL interface: E112, salt-bridges to R295 of PARL, E113 binds to a POPE lipid, and E117 interacts with K114 of PINK1 and N238 of PARL (Figure 6d). In the Romo1 import complex, these PINK1 residues are predicted to also form important interactions at the entrance of the protein-channel: E112 with R6 of Tim17, E113 with R18 of Romo1, and E117 with E127 of Tim17 *via* K114 of PINK1 (Figure 6e). In this context isoleucine 111 of PINK1 forms stable hydrophobic integrations with a pair of methionine residues in Romo1 (Figure 6f). Presumably, this helps further stabilise this region in the channel. Perturbation of the interactions, with the 3 glutamates and isoleucine, would limit PINK1 entry into the Romo1-associated complex and matrix import; and thereby, in the case of I111S, contribute to onset Parkinson’s disease. Once incorporated into the Romo1-complex the conformation of the PINK1 TMD remains fully α-helical, in contrast to it forming an α/β-hybrid when associated with the PARL-complex.

Taken all together, it is clear that the TMD and the residues that immediately follow, including I111 and the three conserved glutamates, are critical for the stabilisation of both complexes (Figure 4f, 5 and 6). Their contrasting interactions dictated by the presence of PARL or Romo1 determine the structure of the TMD, and thereby enabling cleavage or matrix import as required.

### Residues following the TMD of PINK1 are critical for both cleavage and matrix import

The structure predictions and simulations of the complexes are credible because they recapitulate know structures of their subunits: PINK1, Tim17, Tim23 and Tim44, while PARL is in good agreement with the experimental determined homologue GlpG. Moreover, simulations of the structures also correctly predict the location of phospholipid binding sites on the TIM23-core-complex. They also visualise key expected features, such as: (1) the juxtapositioning of the PINK1 cleavage site and the PARL active site; (2) the constricted membrane and water filled cavity at the protease site; (3) the α/β-hybrid TMD, and (4) the protein-channel through the inner-membrane.

To further validate the models and provide additional insights into the mechanism of PINK1 cleavage and import, we carried out import assays of the variant, wherein the 3 highly conserved glutamates were substituted by alanine (E112A, E113A and E117A; 3EA), or the version associated with early onset Parkinson’s Disease (I111S). From the modelling and simulation data, we hypothesised that these substitutions should destabilise both PARL and Romo1 associated complexes, thereby affecting PINK1 cleavage and matrix import.

Regarding cleavage by PARL, this has already been shown to be diminished in both I111S and 3EA ^52,53^. To determine their impact on mitochondrial import, we applied our MitoLuc matrix import assay. Again, their 11S activation properties were measured (Figure S12 and S13) and used to scale the luminescent readout corresponding to import. The results show that the 3EA variant has a major impact on import into the matrix. Only tiny amounts of the N-(LDD) and C-terminal (DDL) sections of PINK entered the matrix, while the central section (DLD), just prior to the kinase domain, was affected to a lesser degree, being reduced by about half (Figure 7a-c). This pattern was reproduced, but less pronounced, for IMS import; the N- and C-terminal reporters were reduced by ∼50%, while the central section was unaffected (Figure 7d-f). In respect of IMS import, the two rate determining steps of transport across the OM were unaffected by the 3EA amino acid substitutions (Figure 7g). This confirms that the effect is a not a consequence of altered interactions with the TOM complex. The lower yield of IMS entry is presumably a result of restricted transport further downstream, through the TIM23-complex. Thus, both cleavage ^52^ and transport across the inner-membrane (but not the outer-membrane) of the 3EA variant are reduced.

**Figure 7:**
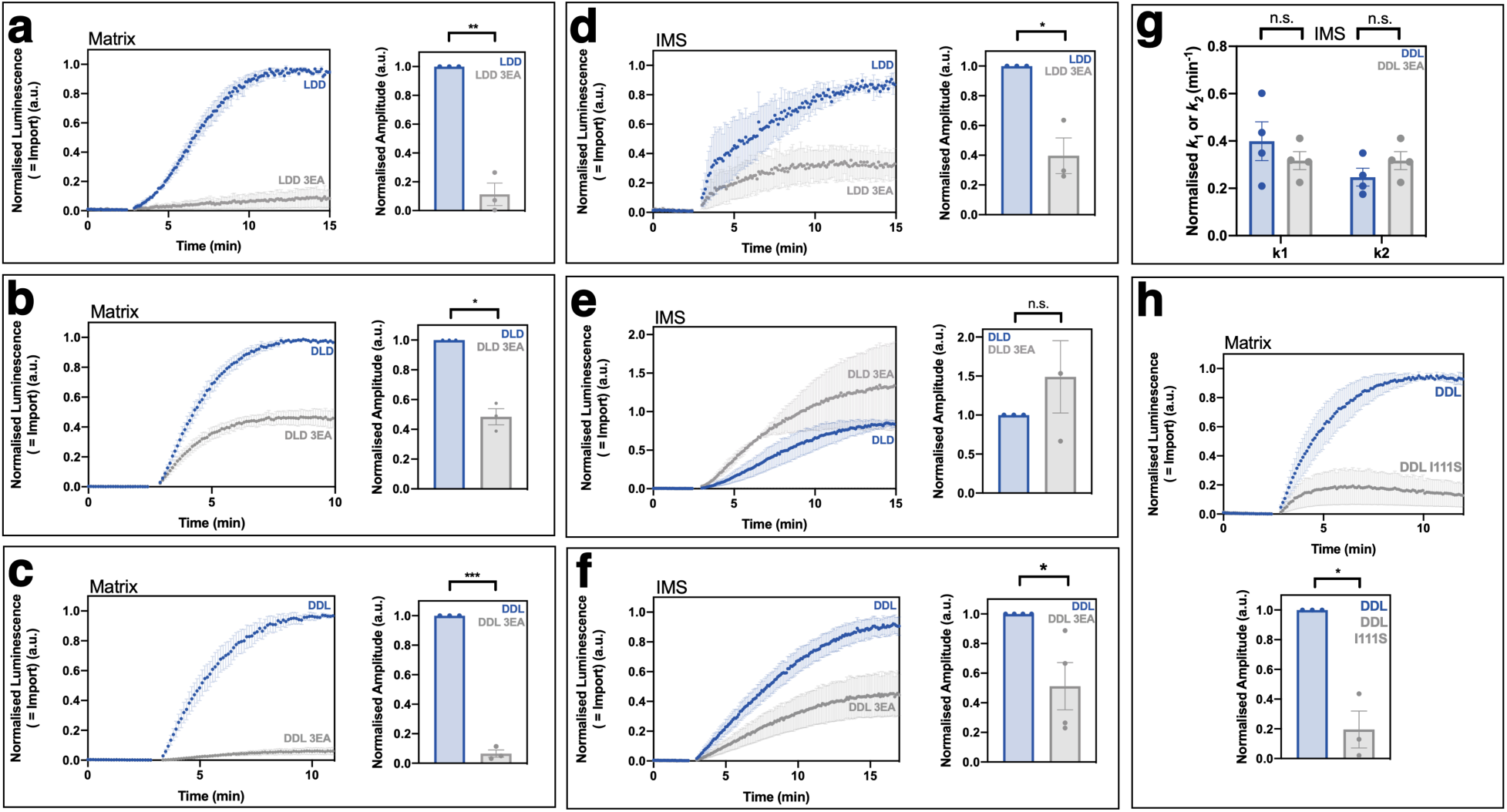
The 3 conserved glutamates and I111S of PINK1 are critical for matrix import. Import traces for import of PINK1 versus a variant wherein the 3 conserved glutamates are substituted with alanine (3EA). Data shown for entry of PINK1 into the IMS for the constucts LDD (**a**), DLD (**b**) and DDL (**c**); repeated for entry into the matrix (**d-f**). **(g)** Comparison of the rate constants determined by fitting the MitoLuc data for IMS import (PINK1 DDL and PINK1(3EA) DDL) to a two-step kinetic model. A one-way ANOVA with Tukey’s post hoc test was used to determine the significance of the differences of k_1_ and k_2_. **(h)** MitoLuc import trace and associated amplitude data for the PD-linked PINK1 variant I111S. Error bars show SEM of the data (N=3 biological repeats) with each biological repeat calculated from three technical repeats. A paired t-test was used to compare the normalised amplitude data.

Substitution of I111S also affects PINK1 import. Complete matrix entry, measured by import of the C-terminus (PINK1-DDL), is reduced (Figure 7h), though less so compared to 3EA. Therefore, I111S and 3EA act similarly, diminishing both PINK1 cleavage ^52,53^ and matrix import. This is consistent with the simulated models highlighting potential key interactions of these residues of PINK1 within complexes mediating both proteolysis (with PARL) and inner-membrane transport (with Romo1).

### Subverting PINK1 TMD structural plasticity (α/β-hybrid formation) favours matrix import

The structural modelling and simulations suggest that the TMD of PINK1 can form either an α-helix or an α/β-hybrid, respectively in association with Romo1 (for import) or PARL (for cleavage). Therefore, we hypothesised that stabilisation of this region as an α-helix (destabilisation of the β-strand) should promote the import pathway. To test this, we created a PINK1 DDL variant (Figure S14) wherein the helix breaking glycine residues (Figure 1a, 4f and S1) were substituted with alanines, which exhibit the highest propensity for promoting α-helix formation, without changing side chains dramatically ^54–57^. The resulting PINK1(ALALAL) variant should be far less likely to form a β-strand with PARL and thereby favour matrix import. As anticipated, the ALALAL variant demonstrates higher import than the native GLGLGL PINK1 (Figure 8), strongly suggesting that the ability to switch from an α-helix to a β-strand is important for selecting between cleavage and import pathways.

**Figure 8:**
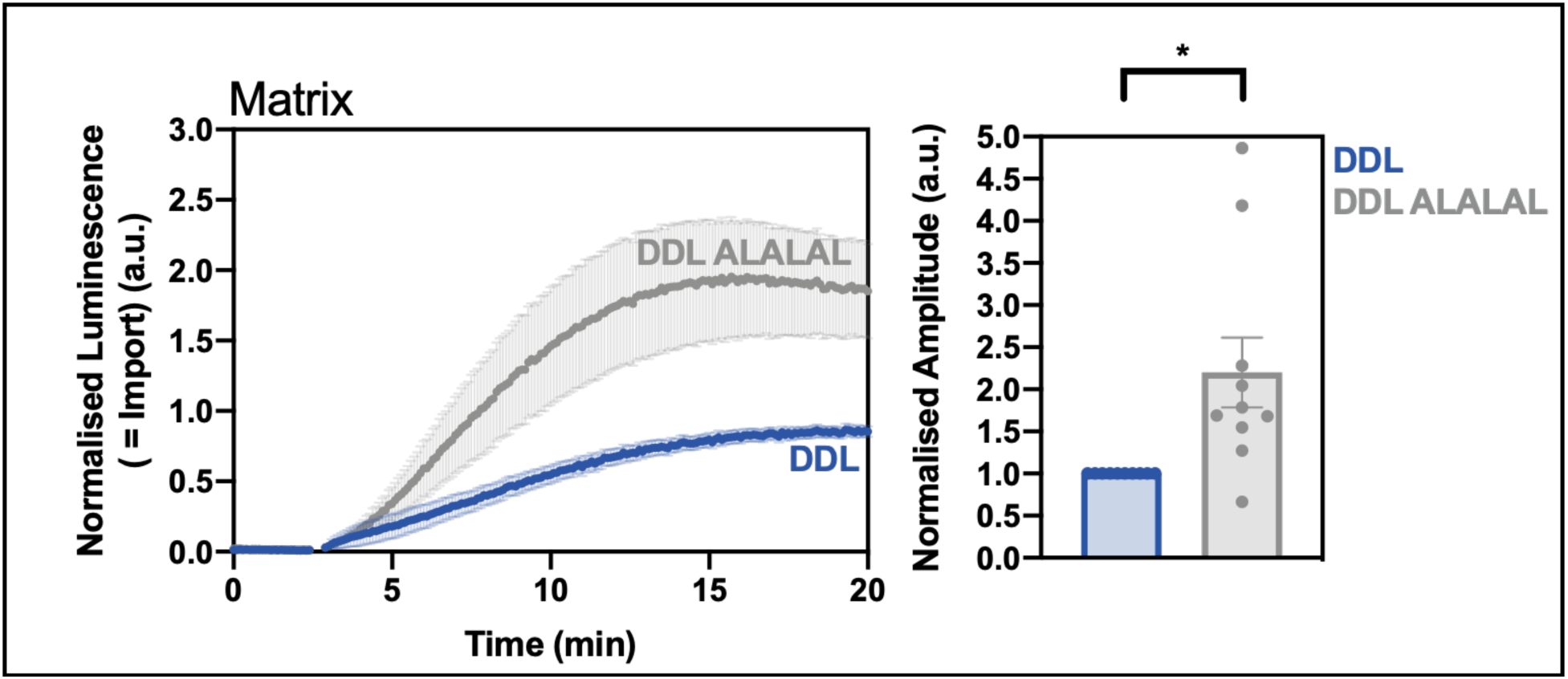
Promoting Helix Formation within PINK1’s TMD Increases its Import into the Matrix. **(a)** MitoLuc analysis of the import into the matrix of PINK1 DDL compared to the equivalent construct with GLGLGL substituted for ALALAL. Normalized amplitudes were calculated and plotted in **(b)**. Error bars represent SEM and data an N=10 biological repeats where three technical repeats were performed for each biological repeat. A paired t-test was used to confirm significance of the amplitude data.

## Discussion

The successful production and purification of full-length human PINK1, developed for this study, has provided new opportunities for its biochemical analysis. Its availability, combined with our recently established in-cell MitoLuc assay, has enabled the characterisation of PINK1 mitochondrial import. The results, together with structural predictions and molecular dynamics simulations, have inspired a re-evaluation of PINK1 processing and transport.

### An extended model for the mitochondrial import behaviour of PINK1

The initial stages of the new model follow those of the canonical version (Figure 1b). which we have been able to elaborate. We show that only the TMD of PINK1 is essential for targeting (this study), and that precursor passage into the IMS occurs in 2 distinct steps, most likely: (1) association with the TOM-complex (Figure 9–Box 1), and (2) transport across the OM (Figure 9–Box 2). Translocation of precursors across the OM is largely complete before engagement with the TIM23-complex (Figure 9–Box 2) ^7^.

**Figure 9:**
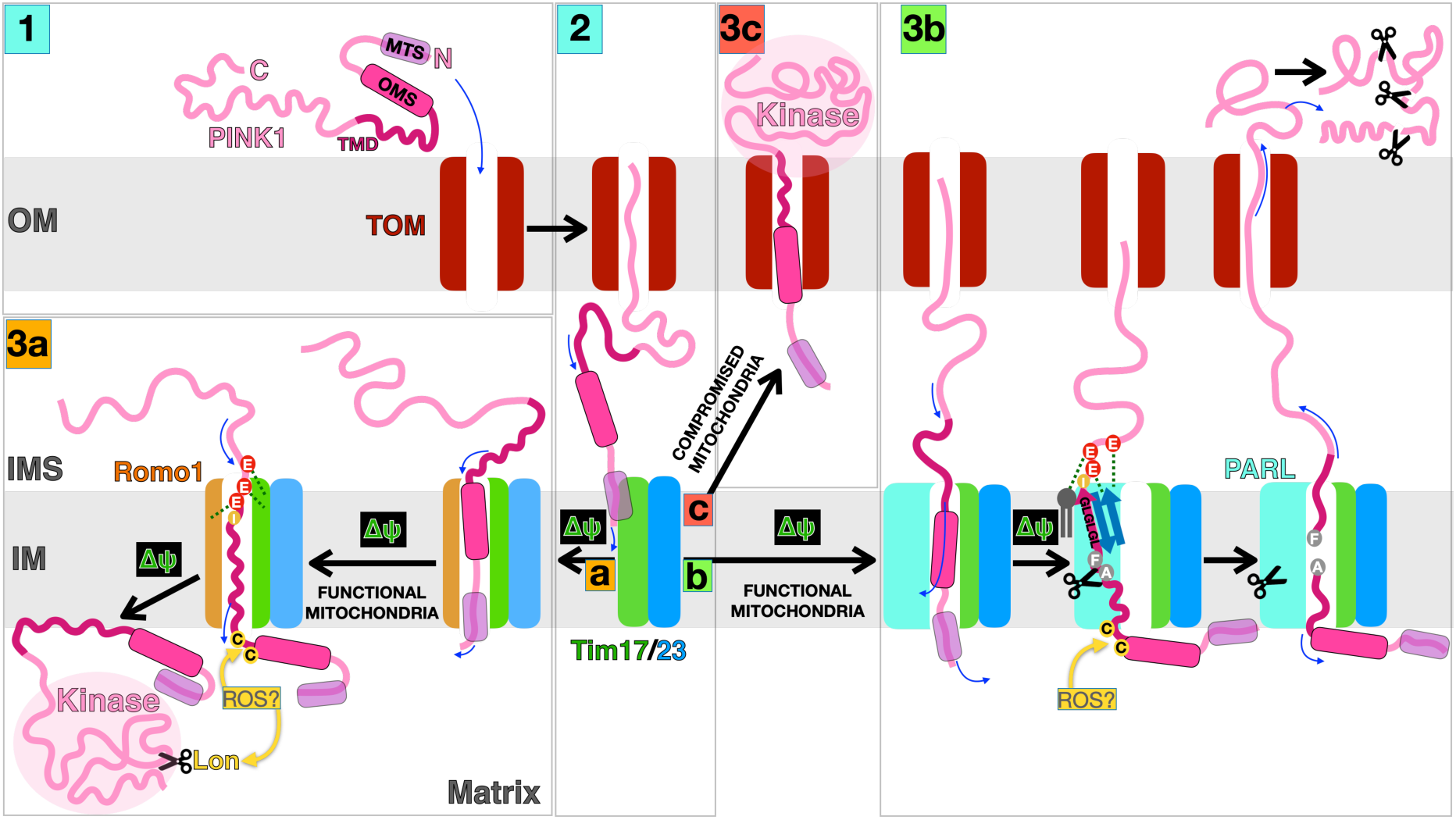
A Revised Model for Mitochondrial Import of PINK1. Schematic overview of the different destinations of PINK1; see text for further details. Key elements of PINK1 are labelled in Box 1; mitochondrial targeting sequence (MTS), outer-membrane targeting sequence (OMS), trans-membrane domain (TMD), with consistent colouring throughout; blue arrows indicate the direction of protein translocation. According to the current model PINK1 is transported through the TOM-complex into the IMS to make contact with the TIM23-complex (Box 1→2). In functional mitochondria, generating a membrane potential (Δψ), PINK1-Tim17-23 recruit either Romo1 (Box 2a) or PARL (Box 2b)–both completing a channel for translocation of the N-terminus across the membrane, driven by Δψ. In the former case, PINK1’s TMD forms an α-helix which can readily be transported into the matrix along with the rest of the protein, again driven by Δψ (Box 3a). While, the interaction with PARL promotes transport into the matrix and entry of the TMD into the inner-membrane (Box 3b). Formation of the α/β-hybrid promotes retraction of the TMD from the channel into the proteolytic active site for cleavage between phenylalanine (F) and alanine (A). The respective cleavage produces can be transported in opposite directions into the IMS and matrix. Retro-translocation of the C-terminal fragment back through the TOM-complex results in degradation by the proteosome. When mitochondria become compromised the loss of Δψ mean that PINK1’s N-terminus is not transported and the full-length protein is then trapped at the outer-membrane (Box 3c), leading to dimirisation, kinase activation and Parkin recruitment. The 3 conserved glutamates (E) are shown making important, but distinct, interactions with both Romo1 and PARL associated complexes. Between the 3Es and the TMD is an isoleucine (I), which when substituted with serine (I111S) brings about early onset Parkinson’s disease. ROS may regulate this process by promoting the Lon-dependent degradation of matrix PINK1 (Box 3a) and by the oxidation of activated thiols between PINK1’s OMS and TMD (Box 3a, 3b).

Incursion of PINK1 into the inner-membrane requires the membrane potential ^58^ and proceeds at the interface between Tim17 and exclusively either: the Mgr2-like protein Romo-1 (Figure 9–Box 2a→3a) or PARL (Figure 9–Box 2b→3b). The former resulting in matrix import of full-length PINK1 by the classical TIM23 pathway. For the latter, active engagement of PINK1 with PARL is promoted by inter-protein β-augmentation, and further stabilised by salt bridges of the conserved PINK1-3Es. The ensuing cleavage brings about retro-translocation of the truncated C-terminal fragment followed by its degradation by the proteasome (Figure 9–Box 3b).

When Δψ is diminished (*e.g.* during mitochondrial stress, or artificially with CCCP) PINK1 fails to fully penetrate the inner-membrane, and therefore is neither transported to the matrix, nor is it cleaved by PARL; the latter as previously noted ^58^. Instead, the full-length protein is prone to retro-translocation (Figure 9–Box 2c→3c) and stabilisation at the outer-membrane, which leads to dimerisation, self-phosphorylation, PARKIN recruitment and ultimately mitophagy. This eventuality is also promoted by the replacement of the three key negative charges (3EA), which appears to be due destabilisation of the interaction between PINK1 and PARL (Figure 9–Box 3b), as well as with Tim17/Romo1 (Figure 9– Box 3a). The predicted destabilisation of both of these interactions, according to the model, would prevent both matrix import and cleavage, and thereby promote outer-membrane localisation of the full-length protein (Figure 9–Box 3c)–as observed ^52^. While, destabilisation of only the PINK1-PARL interaction (with the β-breaker) is compatible with full-length matrix import (Figure 9–Box 3a).

Another contributing factor for PINK1 localisation is ROS, as outer membrane accumulation is known to be dependent on mitochondrial redox status ^59^. Our structural model highlights a potential ‘redox-sensor’: two activated cysteines at the end of PINK1’s TMD poking into the matrix. Perhaps increased ROS (associated with a mis-firing electron transfer chain) will bring about their oxidation, which we speculate could perturb its proteolysis by PARL (Figure 9–Box 3b) and/ or matrix import (Figure 9–Box 3a). In turn, this could trigger surface presentation and mitophagy (Figure 9–Box 2c→3c). Consistent with this, another perturbation in this very region P95A, between the two cysteines prevents cleavage ^23^. Furthermore, ROS is also thought the promote Lon activity for the degradation of PINK1 (Figure 9–Box 3a) ^36^. Therefore, it is conceivable that the action of ROS could promote the presentation of PINK1 at the surface, as well as its elimination in the matrix. Interestingly, variants affecting this putative redox-sensor of PINK1 (C92F and R98W) are also associated with early onset Parkinson’s disease ^25,60^.

### Implications for PINK1 matrix import?

The delivery of full-length PINK1 into the matrix has many interesting implications. Though not widely recognised, matrix import of PINK1 has been reported previously. These observations include entry of the kinase domain for phosphorylation of Ndufa10, for activation of Complex I (and thereby electron transfer activity) ^35,42^ and also MIC60/mitofilin for the dynamic re-modelling of cristae structure ^61^. As noted above PINK1 is also degraded by the Lon protease, which is located in the matrix ^36^.

In the context of the import process, it is well known that accessory factors associate with the TIM23-core-complex for functional modulation; for example, to facilitate proteins either into or across the inner-mmebrane. Mgr2 is a prime ancillary candidate as a ‘gatekeeper’ to regulate this process ^50^. Indeed, the structural analysis of the yeast Tim17-23-44 complex suggest that the pathway for proteins into the matrix can be formed at the interface between Tim17 and Mgr2 ^12,16,17^. Additional, factors most likely also participate during protein passage into the matrix and for membrane protein insertion.

The association of the TIM23-complex with another accessory factor, PARL, is presumably required for PINK1 cleavage, though not much is known about it. Our structural model of the complex presents us with new insights into this presumption and its consequences. One important feature is that the interaction of the core TIM23-complex with Romo1 and PARL are mutually exclusive. The former results in the encapsulation of the PINK1 TMD as an α-helix within the protein-channel, formed between Tim17 and Romo1, whereas the presence of PARL results in the withdrawal of PINK1’s TMD from the protein-channel for remodelling (α/β-hybrid formation) and cleavage. This dynamic interplay between PINK1 and PARL is compatible with their role in the regulation of mitophagy in response to mitochondrial bioenergetic failure (Figure 9–switching from Box 2→3b to 3c). Whilst, the alternative interaction with Romo1, for matrix entry of PINK1 including its kinase domain, could help regulate mitochondrial structure and activity, independently of mitophagy.

### An alternative protein-channel?

Interestingly, in the alternative complexes the protein-channel formed at the Tim17/Romo1 and Tim17/PARL interfaces seem to be similar. While, in the latter case the channel is unoccupied by PINK, there may be circumstances when this channel is utilised for transport through the inner-membrane. The formation of a protein-channel at the interface with a rhomboid ‘half-channel’ has been noted before. The retro-translocation of proteins from the endoplasmic reticiculum (ER) lumen back to the cytosol for ER-associated degradation (ERAD) has been proposed to occur between Hrd1 and the rhomboid-like Der1 ^62^. By analogy, the channel between Tim17 and PARL could be required for the entry of PINK1 into the inner-membrane, as well as for its retro-translocation back to the cytosol for degradation.

The prediction that the rhomboid PARL is able to contribute in the formation of a hydrated protein-channel together with Tim17, may help solve the problem identified earlier–that the Mgr2 (yeast) and Romo 1 (human) counterparts are not essential proteins. Perhaps, a number of other membrane proteins (like PARL, but non-proteolytic) are also able to encapsulate a protein-channel, and thereby act, when required, as an alternative to Mgr2/ Romo1.

### Discrimination between alternative matrix and outer-membrane destinations of PINK1

The extended model raises an important question about the fate of PINK1: how is delivery to the outer surface or matrix determined? It seems likely that the TMD plays a key role. Its alternative interactions with Romo1 or PARL, and the respective adoption of the GLGLGL-motif as an α-helix or α/β-hybrid, seems to be the deciding factor–for matrix import, or cleavage and retro-translocation. Indeed, when we stabilise the helix by substitution with ALALAL then import is enhanced; due to failure of active PINK1 engagement by PARL (Figure 9–Box 2b⇥3b) resulting in a shift to matrix import (Figure 9–Box 2a⟶3a).

Interestingly, this α→β transition is reminiscent of the TMD substrates of other intra-membrane proteases. Indeed, it is known that rhomboid specificity is dependent on helix breaking residues within the TMD ^63^. Structures of the γ-secretase complex with its clients Notch and the amyloid precursor protein (APP) also show the induction of an α/β-hybrid TMD within the active site of the protease ^48,64^. The similarity of this arrangement is striking; they all involve β-augmentation of the substrate and protease, wherein the cleavage site is between the α-helix and β-strand of the TMD.

The relatively weak hydrophobicity of PINK1’s TMD might account for the destabilisation of the TMD α-helix critical for its structural plasticity, and suit both PARL and Romo1 promoted alternative destinies. The more hydrophilic TMD α-helical state resists stalling in the inner-membrane and is free to proceed into the matrix, while the α/β-hybrid can easily be incorporated into the water rich constriction of the membrane harbouring the active site of PARL. Interestingly, this is also a feature of other rhomboid proteases.

If the TMD is a conformational switch, goverened by distinct interactions between PINK1 and PARL or Romo1, for cleavage or matrix import, then the question remains: what regulates the incorporation of PINK1 into the respective complexes (Figure 9–box 3a or 3b)? Perhaps the putative redox-sensor is important for the modulation of the TMD conformation by ROS, as well as other accessory factors of the TIM23-complex. Alternatively, the loss of membrane potential may have a contrasting effect on the incorporation of PINK1 into the the PARL or Romo1 containing complexes. In which case, varing degrees of mitochondrial stress could impact the balance between kinase degradation, activation at the outer-membrane or in the matrix (Figure 9–box 3a-c).

### Impact of PINK1 transport for human health

It is well known that perturbation of the normal functioning of PINK1 causes human disease. Several mutations of *pink1* have been implicated in early onset Parkinson’s disease ^25^. They result in amino acid substitutions in both the N-terminal region and the kinase domain, affecting mitochondrial transport, cleavage, kinase activity and thereby regulation of mitophagy. The results presented here suggest that the explanation for some of the associated phenotypes could be more nuanced. First of all, the loss of kinase activity will affect mitophagy independent activities of PINK1 within the matrix. Secondly, changes of the N-terminal region could potentially affect the conformational switch within PINK1’s TMD, upsetting the balance between matrix and outer membrane regulatory activity. One notable example being I111S, linked to Parkinson’s disease, previously shown to affect the PINK1-PARL interaction ^53,65^, and which we also demonstrate reduces matrix import. Perhaps then, loss of PINK1 matrix function also contributes to the early onset Parkinson’s disease.

In general, the dynamic properties of PINK1’s TMD are most likely critical for PINK1’s targeting, processing, distribution and activation. Therefore, amino acid substitutions in and around this region could bring about an imbalance of PINK1s localisation and activity, and thereby cause disease. For instance, variants affecting the putative redox-sensor, and potential dynamic response to ROS, at the matrix end of the TMD of PINK1 are also associated with early onset Parkinson’s disease ^25,60^.

Clearly, more work needs to be done to better understand the cues for PINK1 distribution and degradation and their consequences, and how they are affected by the various *pink1* mutants. With this new framework, and further analysis, it is conceivable that new strategies could be developed for the treatment of specific forms of Parkinson’s disease. For instance, by controlling and rebalancing the alternative destinations of PINK1 to correct the mis-regulation of mitochondrial structure, function and quality control. One way to achieve this could be by small molecule interventions that shift the conformational dynamics of its TMD to promote either inner-membrane transport or cleavage as required.

## Acknowledgements

This work was carried out using the computational facilities of the Advanced Computing Research Centre, University of Bristol - http://www.bristol.ac.uk/acrc/. Additional simulations were run using computational resource from ARCHER and JADE UK National Supercomputing Services, provided by HECBioSim, the UK High End Computing Consortium for Biomolecular Simulation (https://www.hecbiosim.ac.uk), which is supported by the EPSRC (EP/L000253/1).

## Author Contribution

JSL and IC devised the project. JSL and RAC produced and analysed the data. All authors interpreted the data and contributed to the writing of the manuscript. IC secured funding and led the project.

## Declaration

The authors declare no competing interests.

## Data and Materials Availability

All data are available in the main text or supplementary material. AlphaFold2 models are available for download at https://osf.io/xj9ca/ along with PAE and pLDDT plots.

## Supplementary Material

### Materials and Methods

#### Cloning

*In silico* cloning including plasmid, gene strand and primer development/modification was carried out on Snapgene software. The vector used for bacterial expression of PINK1 proteins was pCA528-His-SUMO. The mammalian constructs pLenti6-DEST PINK1-V5 WT and pLenti6-DEST PINK1 V5-KD were gifts from Mark Cookson (Addgene plasmids #13319 and #13320; http://n2t.net/addgene:13319/; https://www.addgene.org/13320/; RRID:Addgene_13319/ 13320) ^1^.

Gibson cloning was performed using NEBuilder HiFi DNA Assembly Master Mix (NEB) as per manufacturer’s instructions. PCR reactions were performed using the Q5 High Fidelity DNA Polymerase (NEB) starting with 1ng template DNA and using all other concentrations as indicated by the manufacturer. Subsequent PCR products were purified using the QIAquick PCR purification kit (Qiagen). Commercial NEB restriction enzymes were used for restriction digests in which the reaction was typically carried out at 37°C for 1hr. Subsequent ligations were performed at 16°C overnight using the NEB T4 DNA ligase. The QIAprep Spin Miniprep Kit (Qiagen) and PureYield Plasmid Maxiprep System (Promega) were used for plasmid preparation. Sequencing of plasmids was performed by Eurofins Genomics. Transformations were carried out as indicated below for a variety of competent cells (BL21 (DE3), XL-1 blue and *⍺*-select).

#### BL21 (DE3) Expression Strain and Transformation

In house BL21 (DE3) chemically competent *E. coli* were originally sourced from NEB, prior to generation of in-house lab stocks and were used for expression of PINK1 LD variants. For transformation 50uL BL21 (DE3) bacteria were incubated with 10-50ng DNA for 30min on ice. Bacteria were then heat-shocked at 42°C for 45 seconds followed by 3min incubation on ice. 750uL SOC media was added to the bacteria that were subsequently recovered at 37°C for 1hr. Transformation mixture was subsequently plated on LB agar supplemented with either 50ug/mL kanamycin or 100ug/mL ampicillin.

#### HEK 293T Cell Culture

HEK 293T cells were maintained in ventilated T-75 flasks in CO_2_ incubators supplying 5% CO_2_ with a maintenance temperature of 37°C. Cells were kept in glucose media (High glucose DMEM (Gibco 11965-092), 10% FBS (Gibco, 10437-028) 1xpenicillin streptomycin (P/S, Sigma)) and passaged at approximately 70% confluency. Passaging was carried out by first washing cells with 1x HBSS followed by detachment in 0.05% Trypsin-EDTA (1x) (Gibco, 25300-062). Glucose media was subsequently added to deactivate trypsin after approximately 1-5min and cells subsequently seeded at the appropriate density for the downstream experiment.

#### Transfection

Cells were grown to approximately 70% confluency and subsequently transfected with 1ug DNA using Lipofectamine 3000 (ThermoFisher Scientific). Cells were subsequently incubated at 37°C for the desired expression window, typically 48-72hr.

#### PINK1 LD Series and Variants Purification

A 100mL PINK1 LD variant transformed BL21 (DE3) bacterial pre-culture supplemented with 50ug/mL kanamycin was grown overnight at 37°C shaking at 200rpm. The following morning pre-culture was used to inoculate 4L kanamycin-supplemented LB culture. Cells were allowed to grow at 37°C for approximately 2hr until O.D. had reached approximately 0.6-0.8. Expression was then induced following addition of 0.1mM isopropyl ß-D-1-thiogalactopyranoside (IPTG) and subsequent growth at 37°C for 3hr. Cells were harvested by centrifugation at 5000 x g for 15min (Sorvall LYNX 6000). Cell pellets were subsequently resuspended in TG buffer (50mM Tris pH 8.0, 8M guanidine-HCl, pH 8.0), left shaking on ice for 20min and then sonicated at 100% amplitude for a cycle of 5×40sec on, 40sec off (Fisherbrand Model 120 Sonic Dismembrator). Cell lysates were clarified by centrifugation at 38,000rpm for 45min at 4°C (Beckman Optima XPN-80) before the soluble fraction was loaded onto a chelating Ni^2+^ Sepharose fast flow column (Cytiva, 17057502) equilibrated in TG buffer. The column was then washed with TG + 30mM imidazole and then TNU buffer (50mM Tris, 500mM NaCl, 6M urea, pH 8.0) + 30mM imidazole. Protein was eluted in TNU buffer + imidazole over a gradient of 1CV from 0-500mM imidazole. Pooled elution fractions were subsequently stepwise diluted to 2M urea in TN buffer (50mM Tris, 500mM NaCl, pH 8.0), BME added to 1mM and then cleavage of the SUMO tag performed by addition of in-house purified Ulp1 for 1hr at room temperature. The cleaved sample was subsequently diluted in TNU to dilute the imidazole concentration to 30mM prior to reloading onto the same in-house packed Ni^2+^ column pre-equilibrated in TNU + 30mM imidazole. The flowthrough containing the cleaved PINK1 precursor was collected and dialyzed overnight in TU buffer (50mM Tris, 6M urea, pH 8.0) at 4°C. The following day the dialyzed sample was loaded onto a 5mL HiTrap SP HP column (GE Healthcare) equilibrated in TN(50)U (50mM Tris, 50mM NaCl, 6M urea, pH 8.0). The column was washed in TN(50)U buffer prior to elution over a gradient of 0-500mM NaCl in TNU buffer. Pooled elution fractions were concentrated in a 30K MWCO centrifugal filter (Millipore) at 4000 x g before being aliquoted, snap frozen in liquid nitrogen and stored at -80°C.

#### Purification of ACP1-pep86

BL21 (DE3) cells expressing the ACP1-pep86 precursor were cultured in LB media overnight prior to being sub-cultured into 2xYT media. On reaching OD_600_ = 0.6 cells were induced by addition of IPTG. Cells were harvested 2-3hrs post-expression and lysed via a cell disruptor (Constant Systems Ltd.). Inclusion bodies containing ACP1-pep86 were progressively solubilised into 1xTK buffer (20mM Tris pH 8.0, 50mM KCl, pH 8.0) + 6M urea and then loaded onto a 5mL HisTrap FF column (Cytiva). The column was washed in 1xTK + 6M urea + 50mM imidazole and protein eluted at 300mM imidazole. Pooled elution fractions were loaded onto a 5mL Hi Trap Q HP anion exchange column (Cytiva) and a salt gradient up to 1M KCl applied to elute the bound protein. Desalting was carried out by centrifugation in a 10,000 MWCO spin concentrator and subsequent dilution in 1xTK buffer + 6M urea. An extinction coefficient of 14,440M^−1^cm^−1^ was used to determine protein concentration.

#### Purification of MitoLuc Components

GST-tagged recombinant perfringolysin (rPFO), GST-Dark peptide and His-tagged Su9-EGFP-pep86 were purified as in ^2^.

#### Permeabilised Cell MitoLuc Import Assay

HEK cells were seeded at 600,000 cells/well in 6-well plates and transfected with pXLG3-eqPF670-P2A-Cox8a-11S (matrix 11S) and incubated at 37°C for 48hrs. Alternatively, cells were transfected with pXLG3-eqPF670-P2A-Smac/DIABLO(1-57)-11S (IMS 11S) and incubated at 37°C for 24hrs. Cells were subsequently trypsinized, diluted in normal glucose growth medium and seeded on 96-well plates at a density of 20,000 cells/well. Cells were cultured overnight at 37°C prior to commencement of the MitoLuc import assay. Cells were washed 2x with 1xmannitol respiration buffer pH 7.3 (225mM mannitol, 10mM HEPES, 2.5mM MgCl_2_, 40mM KCl, 2.5mM KH_2_PO_4_, 0.5mM EGTA) and then 100uL MitoLuc import buffer pH 7.3 (1xMR Buffer, 10uM GST-Dark , 0.1% Prionex, 0.1mg/mL creatine kinase, 5mM phosphocreatine, 1mM ATP, 3nM rPFO, 1:400 Nano-Glo luciferase assay substrate (Promega), 5mM succinate, 1uM rotenone) added to each well. A baseline luminescence was read before proteins were injected at 0.9uM final concentration. Subsequent luminescence was monitored on a CLARIOstar Plus plate reader (BMG LabTech). IMS MitoLuc assay data were fit to the same two-step model as in ^3^.

#### Western Blotting

Protein samples to be analysed by western blotting were prepared first for SDS-PAGE by boiling gel samples at 95°C in 1xLDS + 25mM DTT prior to running on a 4-12% BOLT gel (Thermo Fisher Scientific), run at 200V for 25min. Gels were transferred onto 0.45um nitrocellulose blotting membrane (Cytiva) in 1x transfer buffer (0.34M Tris, 0.26M glycine, 0.14M tricine, 2.5mM EDTA) via the semi-dry Pierce Power Station transfer system (Thermofisher Scientific) at 25V, 2.5mAmp for 10min. Nitrocellulose membranes were subsequently blocked in 1xTBS-T + 5% (w/v) milk for 1hr at room temperature before being probed with primary antibody (Beta actin – Sigma-Aldrich A2228 1:10,000 and PINK1 – cell signalling technology 6946 1:500) in 1xTBS-T + 5% (w/v) milk overnight at 4°C. Membranes were washed in 1xTBS-T + 5% milk for 30min and subsequently probed with secondary antibody in TBS-T + 5% milk. Membranes were once again washed for 30min in 1xTBS-T + 5% milk, incubated with SuperSignal^TM^ West Femto reagent and imaged on an Odyssey Fc (LI-COR).

#### Protein model building

Protein models were built using AlphaFold2, running AlphaFold Multimer v3 via ColabFold v1.5.2 ^4–6^. Models were built as per Table 1. The full PINK1/PARL/Tim17/Tim23/Tim44 complex was constructed by overlaying the PARL from the individual models. No structural clashes were observed, suggesting that this complex is both physically and physiologically plausible. Five rounds were run for each prediction, with the best scoring model used for later analysis. Images were made with PyMOL or VMD. All AlphaFold2 models are available for download at https://osf.io/xj9ca/ along with PAE and plDDT plots.

**Supplementary Table 1:** AlphaFold2 models built for this study. All models are of the human proteins unless otherwise specified.

| AlphaFold2 model | Details |
| --- | --- |
| PINK1/PARL | residues 62-581 of UniProt ID Q9BXM7 and residues 93-368 of Q9H300 |
| PARL/Tim17/Tim23/Tim44 | residues 93-368 of Q9H300, residues 1-143 of Q99595 (TIMM17A), residues 66-209 of O14925, and residues 37-452 of O43615 (37-260 trimmed post-hoc due to low pLDDT – with the amphipathic helix of Tim44 (residues 130-191) included in the figures for illustrative purposes only. |
| Tim17/Tim23/Romo1 | residues 1-143 of Q99595, residues 66-209 of O14925, and all residues of P60602. |
| Yeast Tim17/Tim23/Mgr2 | All residues of UniProt ID P39515, P32897, and Q02889 |
| PINK1_TMD/<br>Tim17/Tim23/Romo1 | residues 1-130 of UniProt ID Q9BXM7 joined by a GSGSGSGS to residues 1-143 of Q99595, residues 66-209 of O14925, residues 298-452 of O43615, and all residues of P60602 |

#### Molecular dynamics simulations

The combined AlphaFold2 PINK1/PARL/Tim17/Tim23/Tim44 complex was used to seed MD simulations (with Tim44 trimmed as described in Table 1). The PINK1 loop between residues 180 and 209 was also removed due to low confidence and replaced with a GSGSGSG linker. Alternatively, simulations were built using PARL/Tim17/Tim23PINK1 with PINK1 trimmed to the TMD (residues 90-130) or the PINK1_TMD/Tim17/Tim23/Romo1 complex with the GSGSGSGS linked removed and PINK1 trimmed to the TMD (residues 90-130). Models were built into simulation systems using CHARMM-GUI ^7,8^. Protein atoms were described with the CHARMM36m force field ^9,10^. Side chain pKas were assessed using propKa3.1 ^11^, and all side chain side charge states were set to their default. The proteins were built into membranes comprising 4.5:4.5:1 POPE, POPC, and cardiolipin (tetraoleyl tails). The membranes were solvated with TIP3P waters and neutralised with K^+^ and Cl^−^ to 150 mM. Each system was minimized and equilibrated according the standard CHARMM-GUI protocol, with a 2 ns final equilibration step. Production simulations were run in the NPT ensemble, with temperatures held at 303.5 K using a velocity-rescale thermostat and a coupling constant of 1 ps, and pressure maintained at 1 bar using a semi-isotropic Parrinello-Rahman pressure coupling with a coupling constant of 5 ps ^12,13^. Short range van der Waals and electrostatics were cut-off at 1.2 nm. Simulations were run to 450 ns with 5 repeats for the full complex, or to 300 or 500 ns for the PARL and Romo1 TMD systems respectively, with 3 repeats.

All simulations were run in Gromacs 2020.1 ^14^. Data were analysed using Gromacs tools and VMD ^15^ Lipid interactions were analysed using the PyLipID package ^16^ using cutoffs of 0.35 and 0.5 nm. Plots were made using or Prism 10.

#### 2.12 Statistics

The inbuilt statistical tests on GraphPad Prism 7 were used for statistical analysis. A combination of unpaired and paired t-tests as well as one-way ANOVA with Tukey’s post hoc multiple comparisons was used to determine if a statistical significance was observed. The p-value cut off for significance was <0.05, with significance values graded as follows: n.s. p=>0.05, * p=≤0.05, ** p=≤0.01, *** p=≤0.001 and **** p=≤0.0001.

### SUPPLEMENTARY FIGURES AND LEGENDS

**Figure S1:**
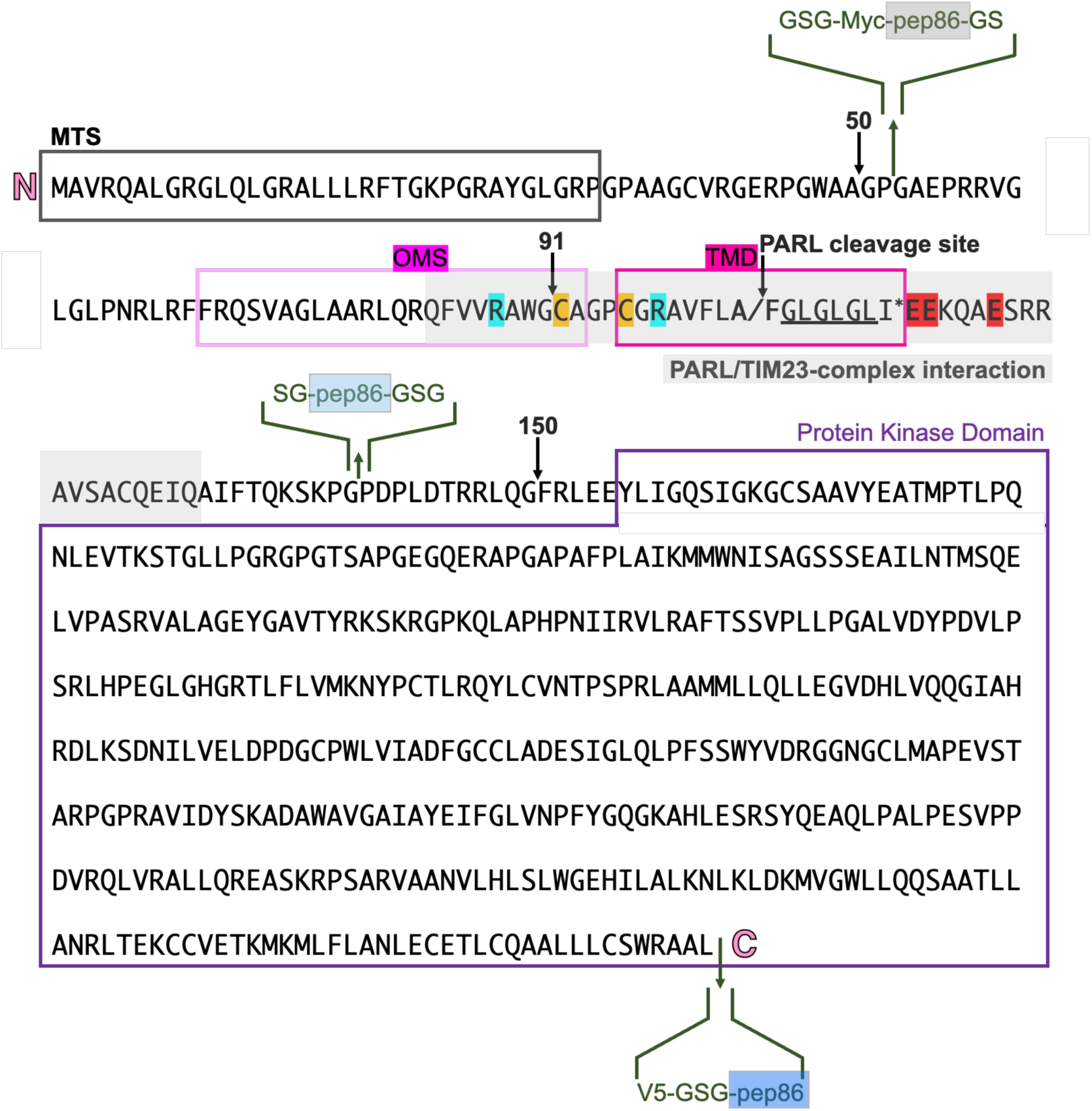
Primary Sequence Depiction and Modification of the Full-length Human PINK1 Protein for MitoLuc Import Studies. The primary sequence of human PINK1 (UniProt: Q9BXM7) is shown annotated with the relevant key domains critical to PINK1 import and function (see text for details). PARL cleavage position and key post-translational modification sites are indicated. Insertion of pep86 and relevant epitope sequences are shown in green, flanked by GSG spacer sequences. Myc and V5 epitope sequences read EQKLISEEDL and IPNPLLGL respectively at the N-terminal to C-terminal insertions.

**Figure S2:**
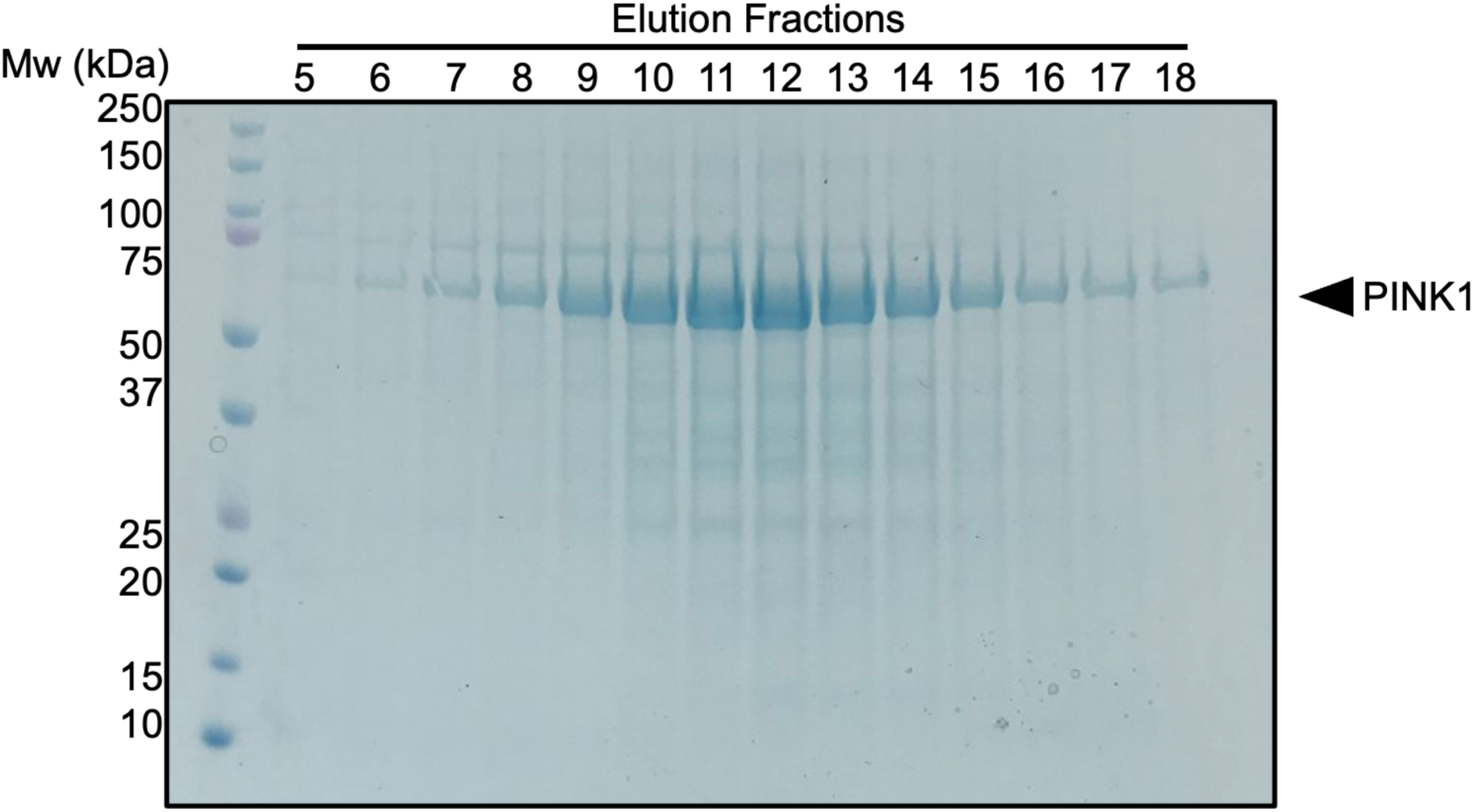
Purification of human PINK1 construct. SDS-PAGE analysis of fractions from cation exchange chromatography, stained by coomasie blue. The visualisation of PINK1 shown here is typical of the purification of all the variants generated for this study.

**Figure S3:**
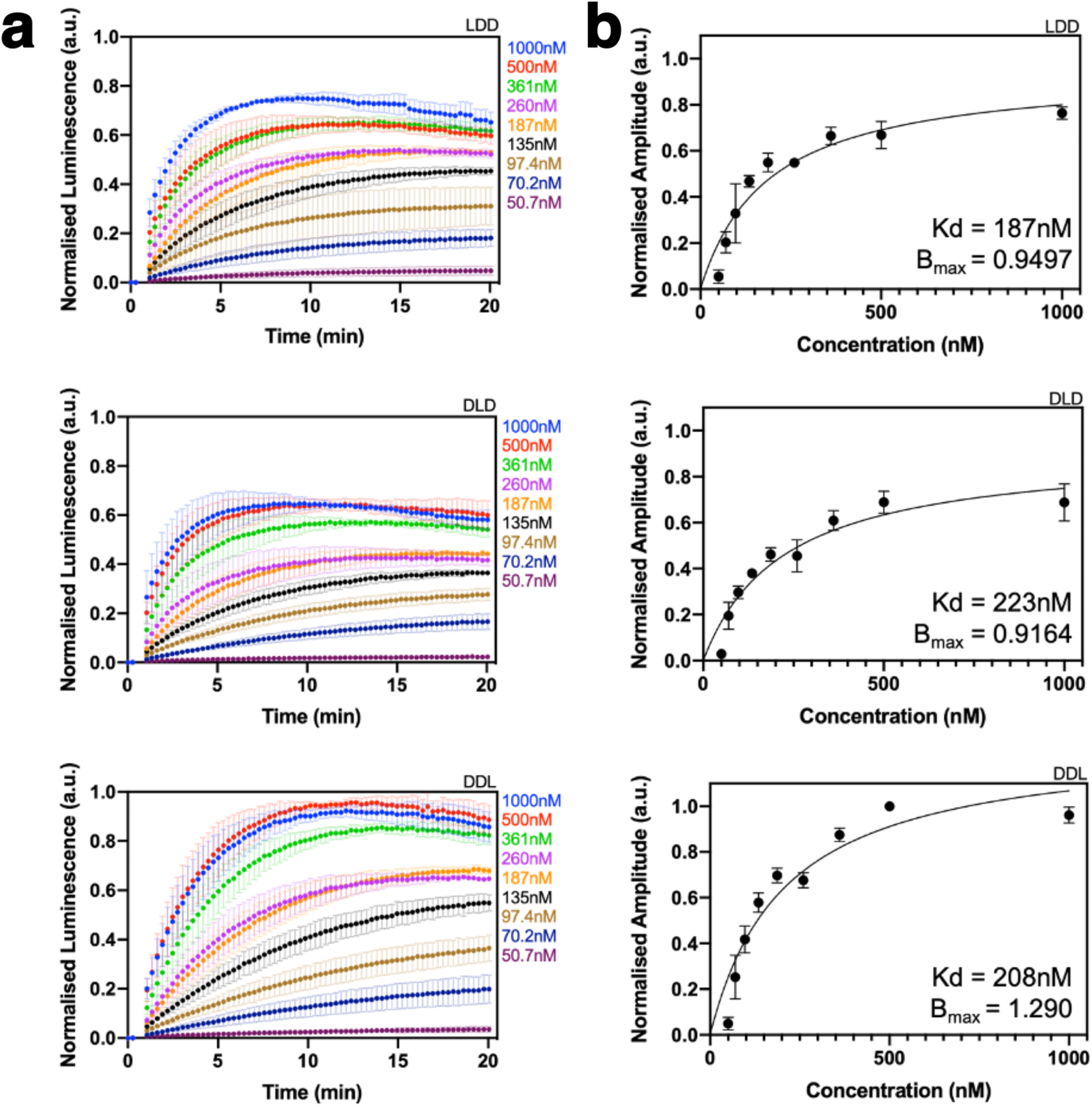
Binding Analysis of the PINK1 LD Series Variants to Isolated 11S. **(a)** Normalised data for PINK1 LDD, DLD and DDL binding to 11S (200pM final concentration) over an LD precursor titration range of 50.7-1000nM. Concentrations and their associated binding curves are indicated by colours. Data represent N=3, error bars for each concentration indicate SEM. **(b)** Normalised amplitude concentration dependence plots for data as in (a). For each LD precursor maximum normalised amplitude was calculated for each concentration and plotted against the corresponding concentration. Data was fit to a one-site binding model (hyperbola). Data represent N=3, error bars for each concentration indicate SEM.

**Figure S4:**
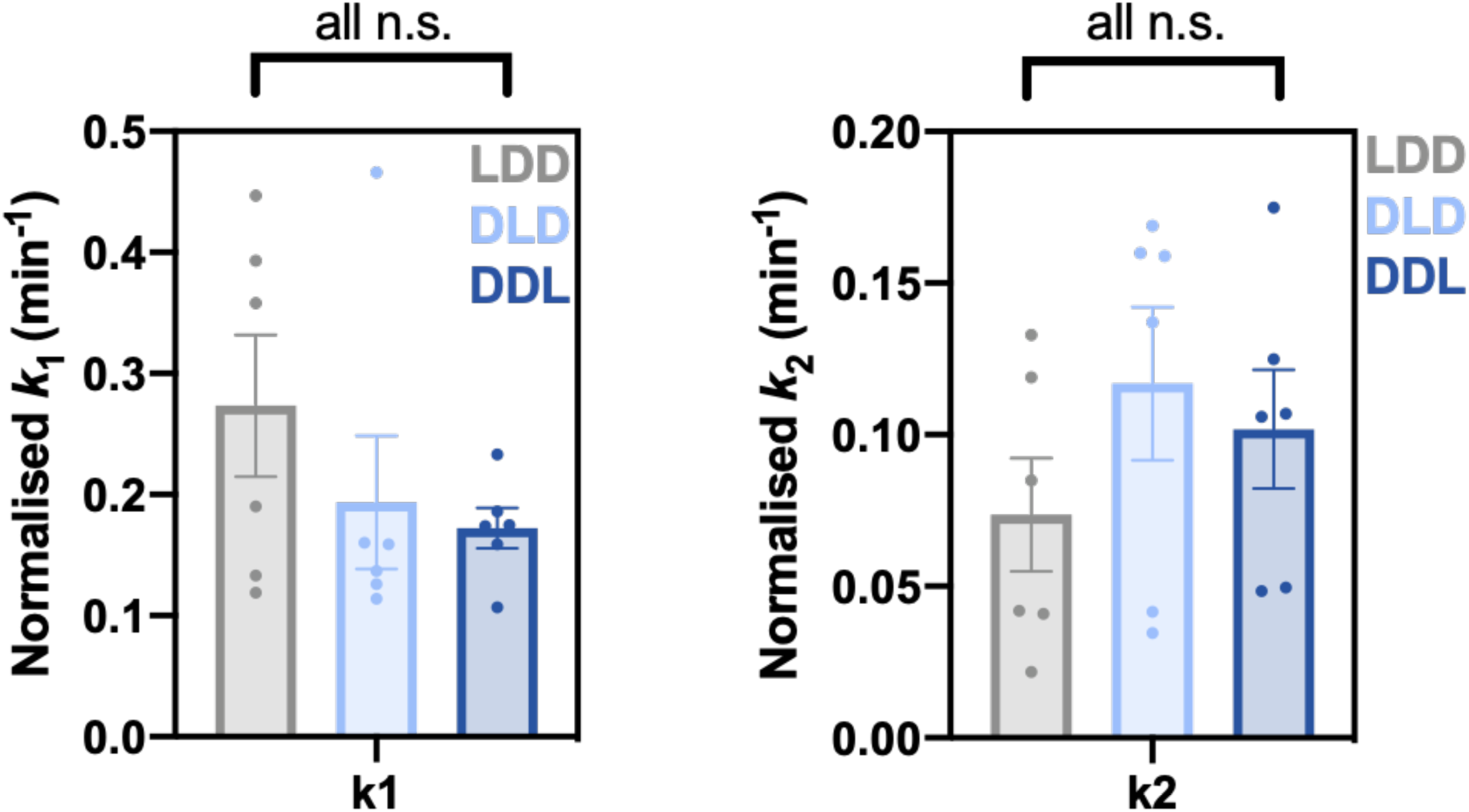
Kinetic Assessment of PINK1 LD Series Import into the IMS. Data for each LD protein was fitted to a two-step model for import and k_1_ **(a)** and k_2_ **(b)** values calculated from the fitted trace. Error bars represent SEM and data are from N=6 biological repeats, where each biological repeat was calculated from three technical repeats. A paired t-test was used to determine significance.

**Figure S5:**
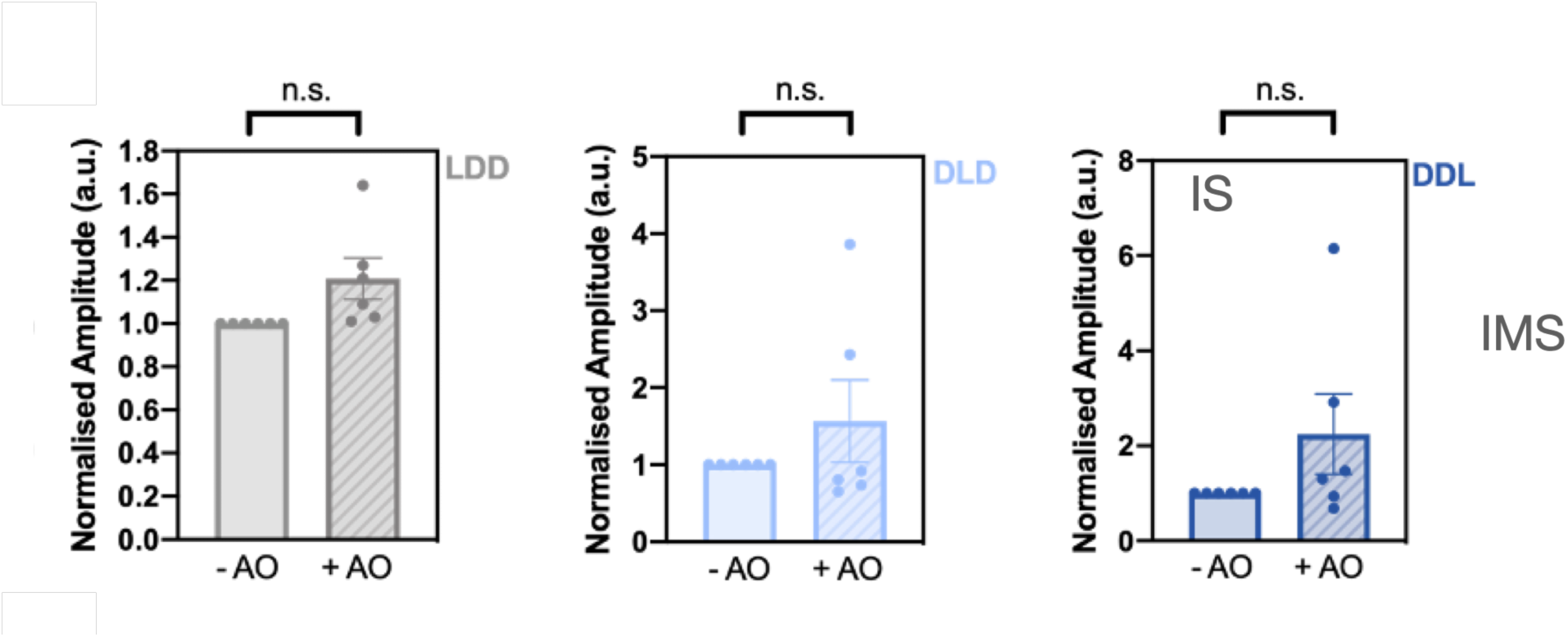
Import of the PINK1 LD Series into the IMS is insensitive to Antimycin A (A) and Oligomycin (O). IMS import data was obtained in the -AO and +AO conditions for each PINK1 LD precursor and the normalised amplitude determined as before. Error bars represent SEM and N=3-6 biological replicates obtained. Amplitude significance was determined using a paired t-test.

**Figure S6:**
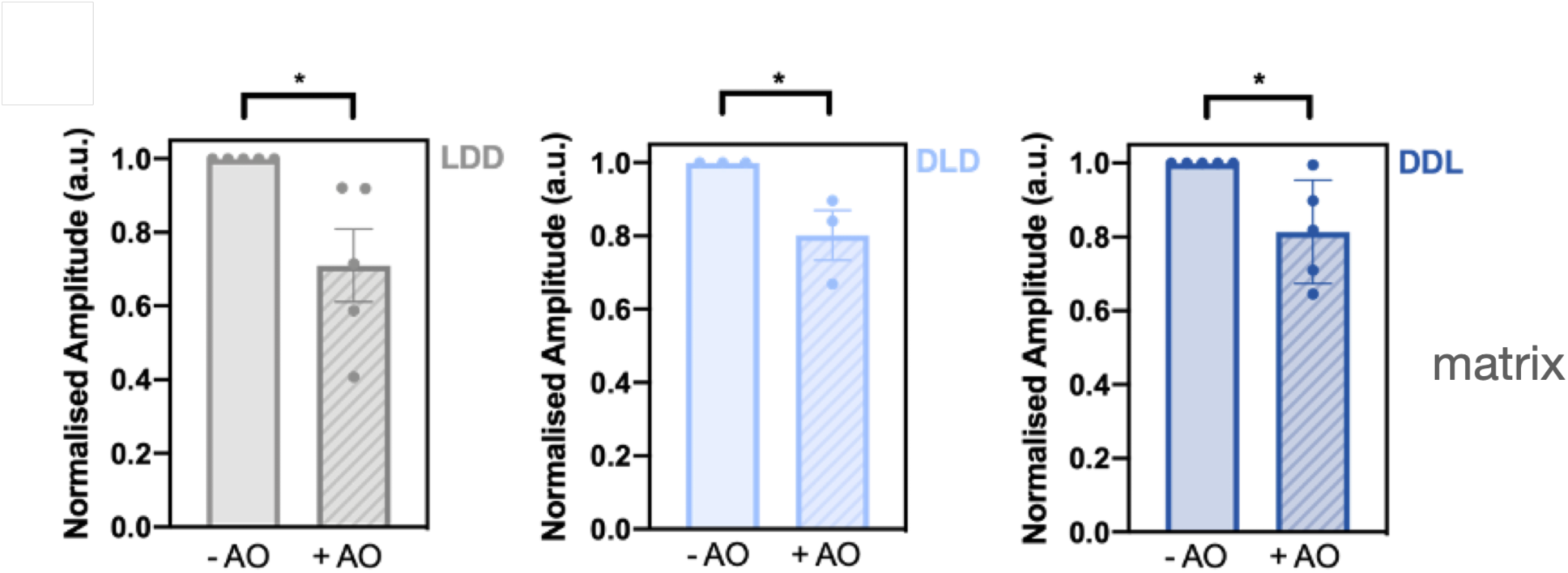
Import of the PINK1 LD Series into the Matrix is is sensitive to Antimycin A and Oligomycin. As described for legend to Figure S5, but for matrix import data.

**Figure S7:**
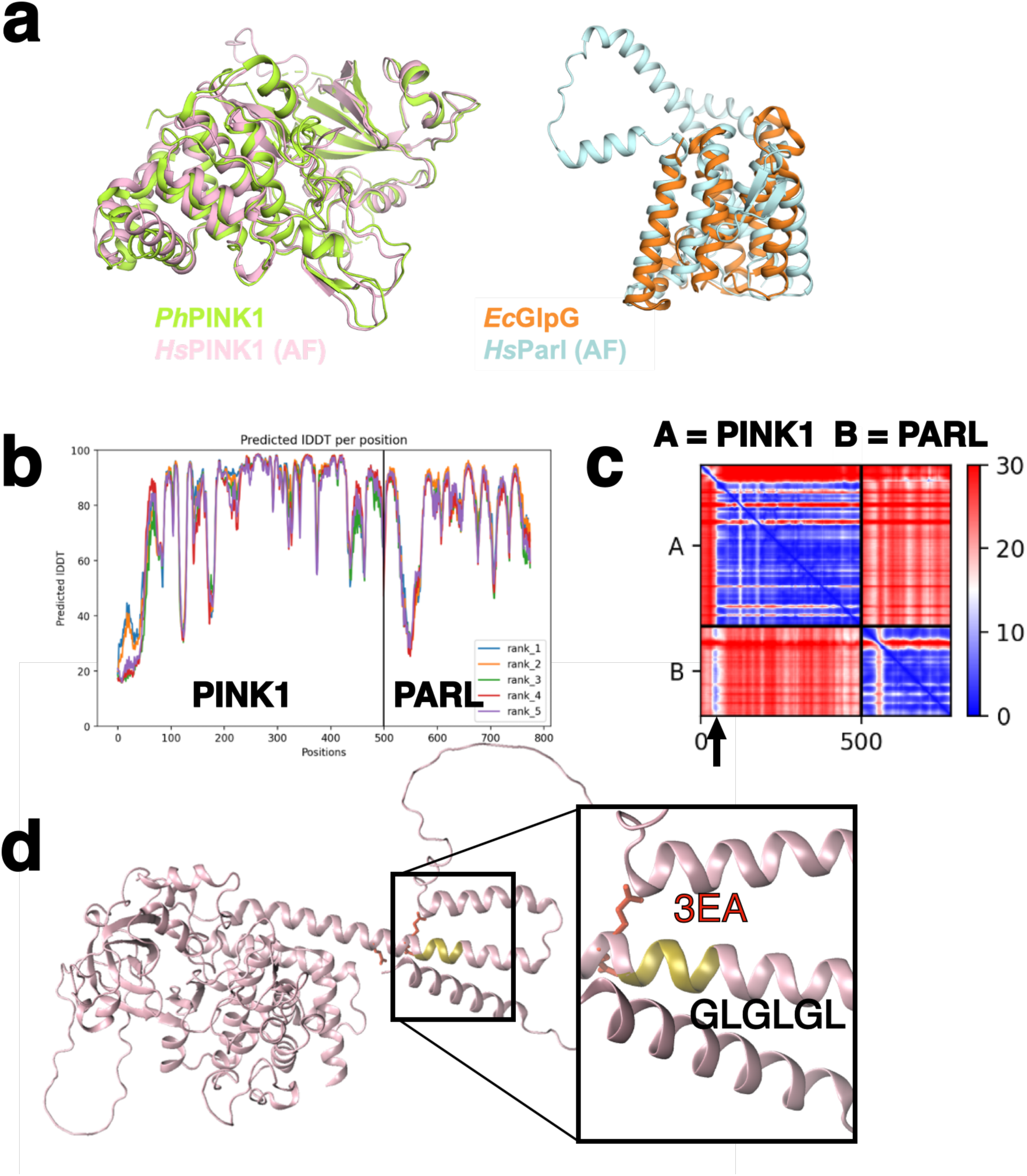
AlphaFold2 modelling of the PINK1/PARL complex. **(a)** Overlays of the modelled human PINK1 with a structure of a homologous PINK1 from *Pediculus humanus corporis* and the modelled PARL aligned to the *E. coli* rhomboid protease GlpG. **(b)** AlphaFold2 pLDDT and for the PINK1-PARL complex model. **(c)** AlphaFold2 PAE scores for the PINK1-PARL complex. An arrow denotes the position of the PINK1-PARL β-strand which corresponds to a high PAE score. **(d)** Human PINK1 from the EBI AlphaFold2 database. The positions of the 3EA and GLGLGL motifs are highlighted on the inset.

**Figure S8:**
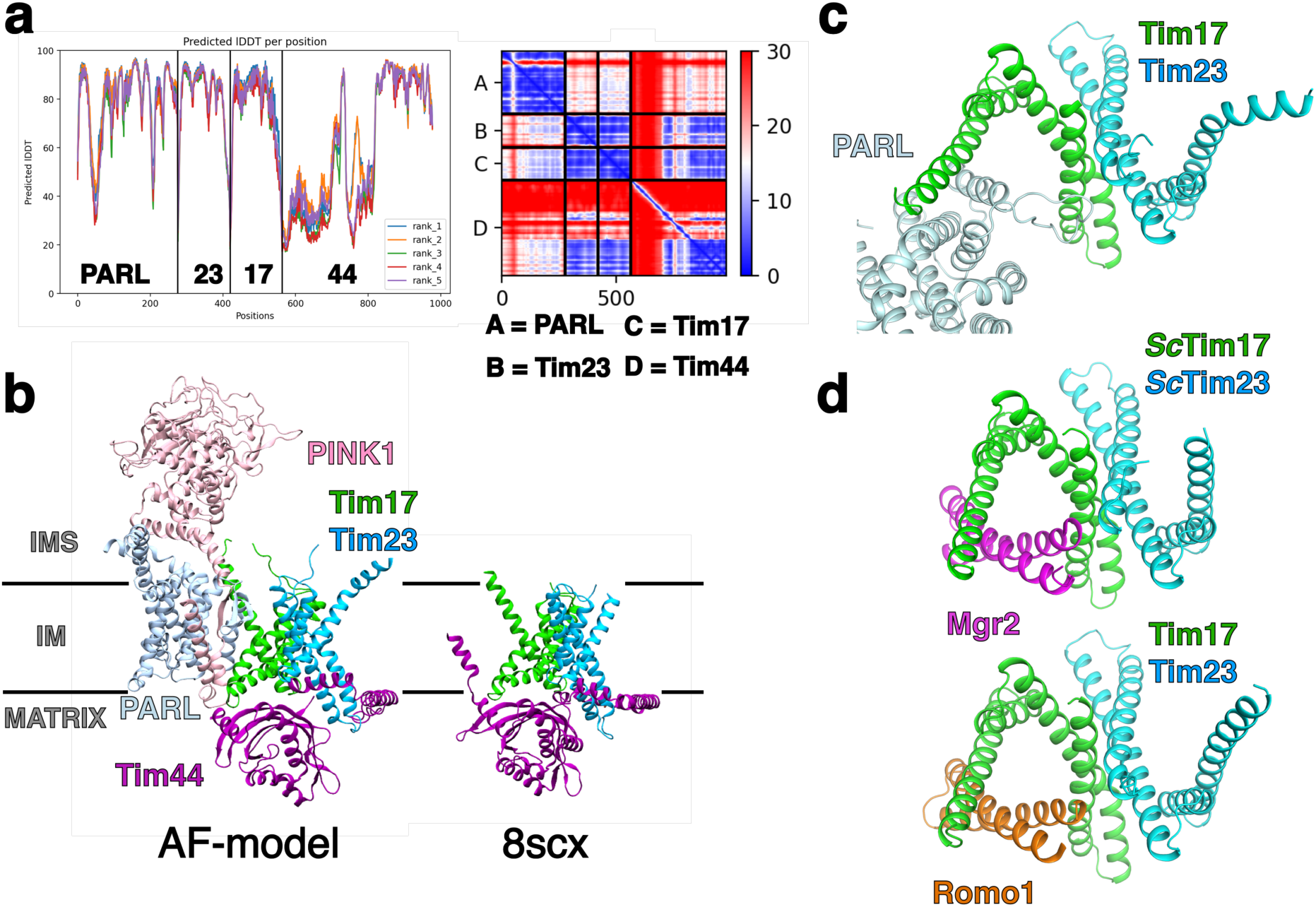
AlphaFold2 modelling of the various complexes containing PINK1, PARL, Tim17, Tim23, TIm44 and Mgr22/Romo1 . **(a)** AlphaFold2 pLDDT and PAE scores for the PARL-Tim17-Tim23-Tim44 complex model. **(b)** View of the hybrid AlphaFold2 PINK1-PARL-Tim17-Tim23-Tim44 model compared to the yeast Tim17-Tim23-Tim44 structure ^17^. The lines indicate the position of the bilayer. **(c)** View of the PARL-Tim17-Tim23 from the same angle as panel. **(d)** Top view of the Tim17-23-Romo1 and yeast (Sc) Tim17-Tim23-Mrg2 AlphaFold2s model showing the presence of the putative protein channel, as in (c).

**Figure S9:**
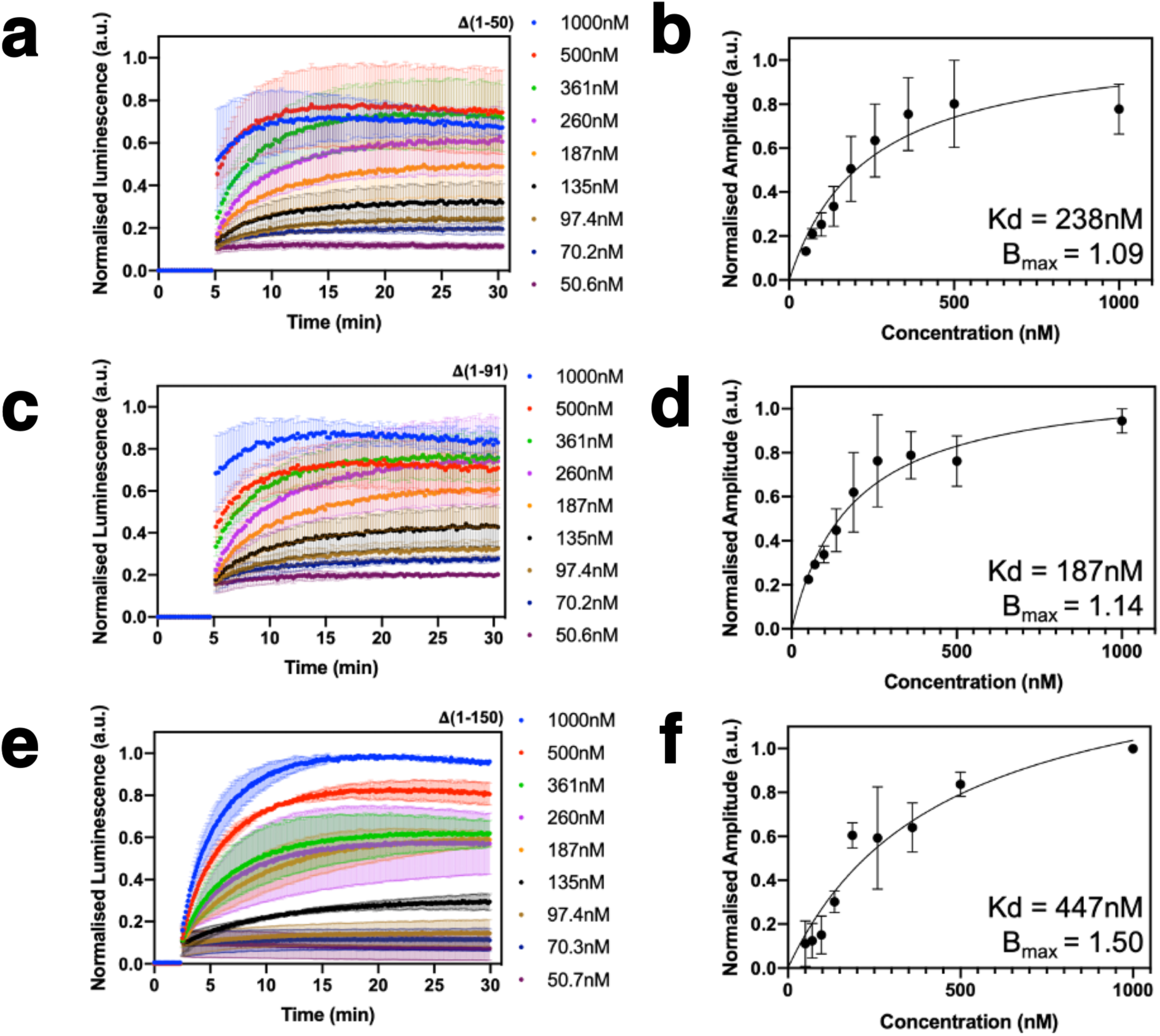
Binding Analysis of the Truncated PINK1 Precursors to Isolated 11S. **(a, c**, and **e)** Titration series of truncated PINK1 precursors was conducted from 50.7nM-1000nM and assayed for binding against 200pM of purified 11S. Maximum amplitudes were determined for each concentration of truncated precursor and plotted in (**b**, **d**, and **f**). Error bars represent SEM and data an N=3 biological repeats.

**Figure S10:**
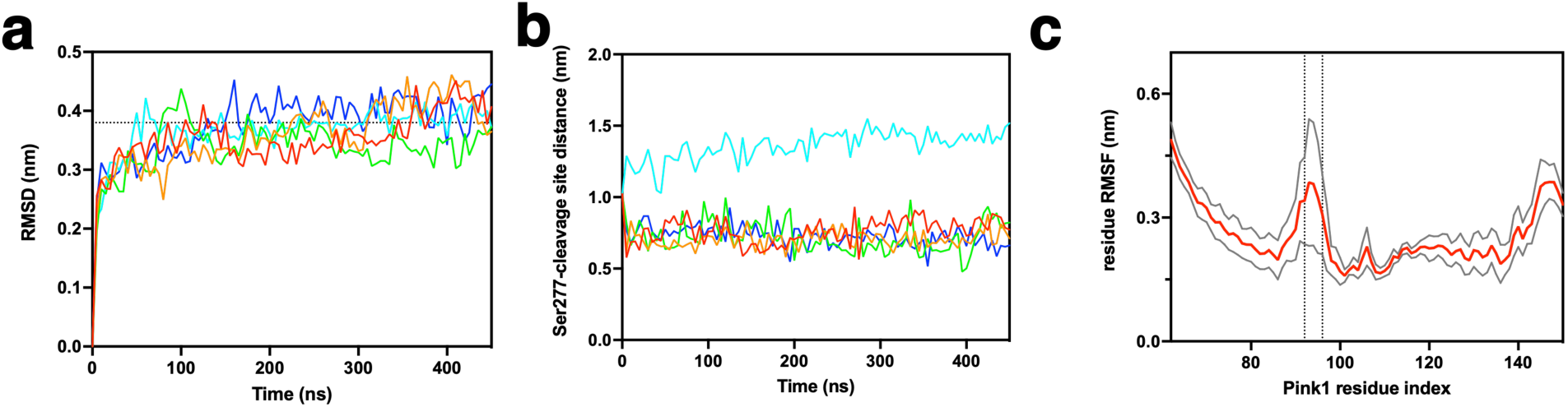
Molecular dynamics supporting data. **(a)** RMSDs of the simulations of the PINK1-PARL-Tim17-Tim23-Tim44 complex (excluding the PINK1 kinase domain). **(b)** Distance plots between Ser277 and the PINK1 cleavage site over the MD simulations. **(c)** RMSF of PINK1 during the PINK1-PARL-Tim17-Tim23-Tim44 MD simulations. Dotted lines denote Cys 92 and 96.

**Figure S11:**
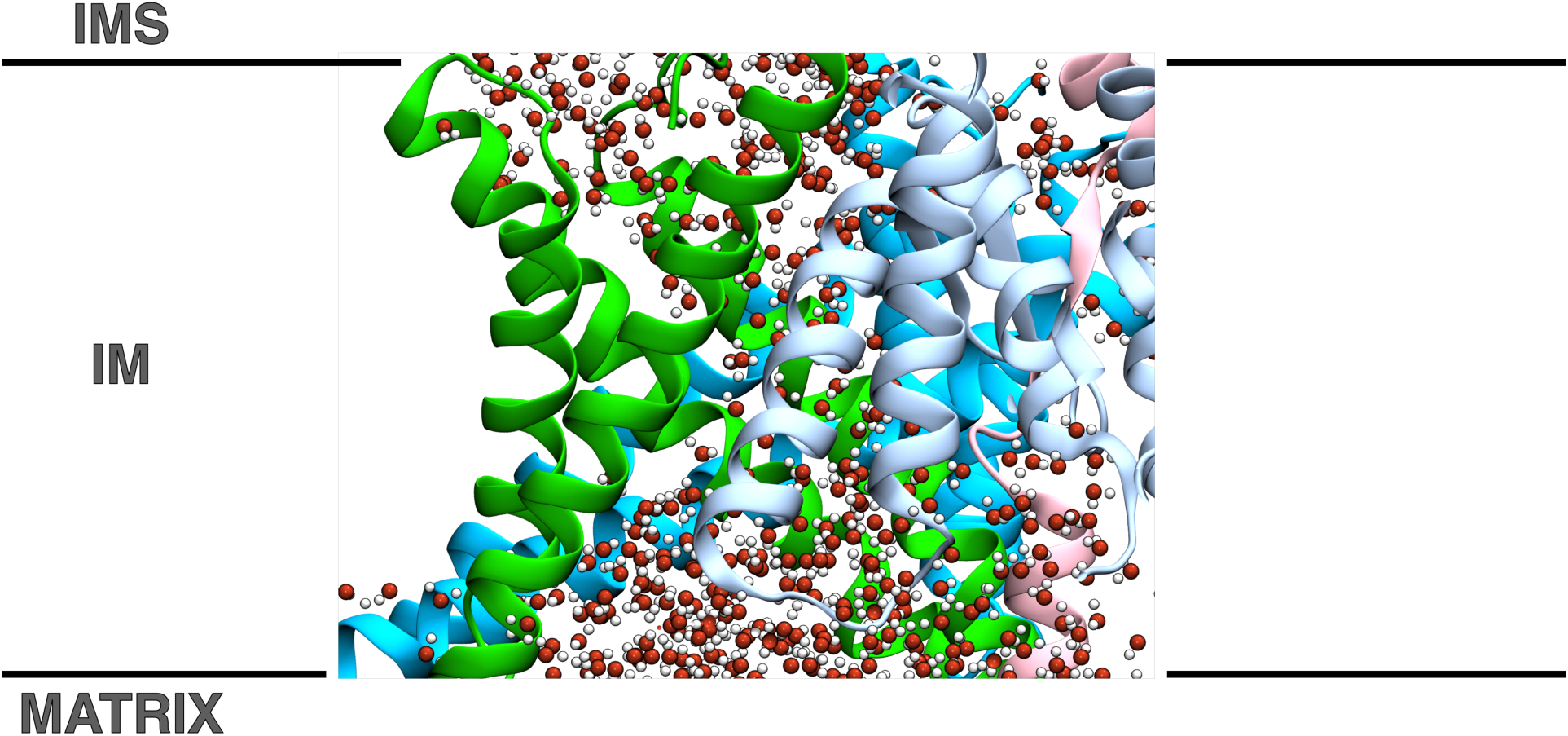
A potential protein-channel at the interface beween Tim17 and PARL. Views of an MD simulation from the side of the membrane, showing Tim17 (green), Tim23 (blue), PARL (cyan) and PINK1 (pink). The location of the water molecules indicate an enclosed solvated channel running through the centre of the complex. The lines indicate the position of the bilayer.

**Figure S12:**
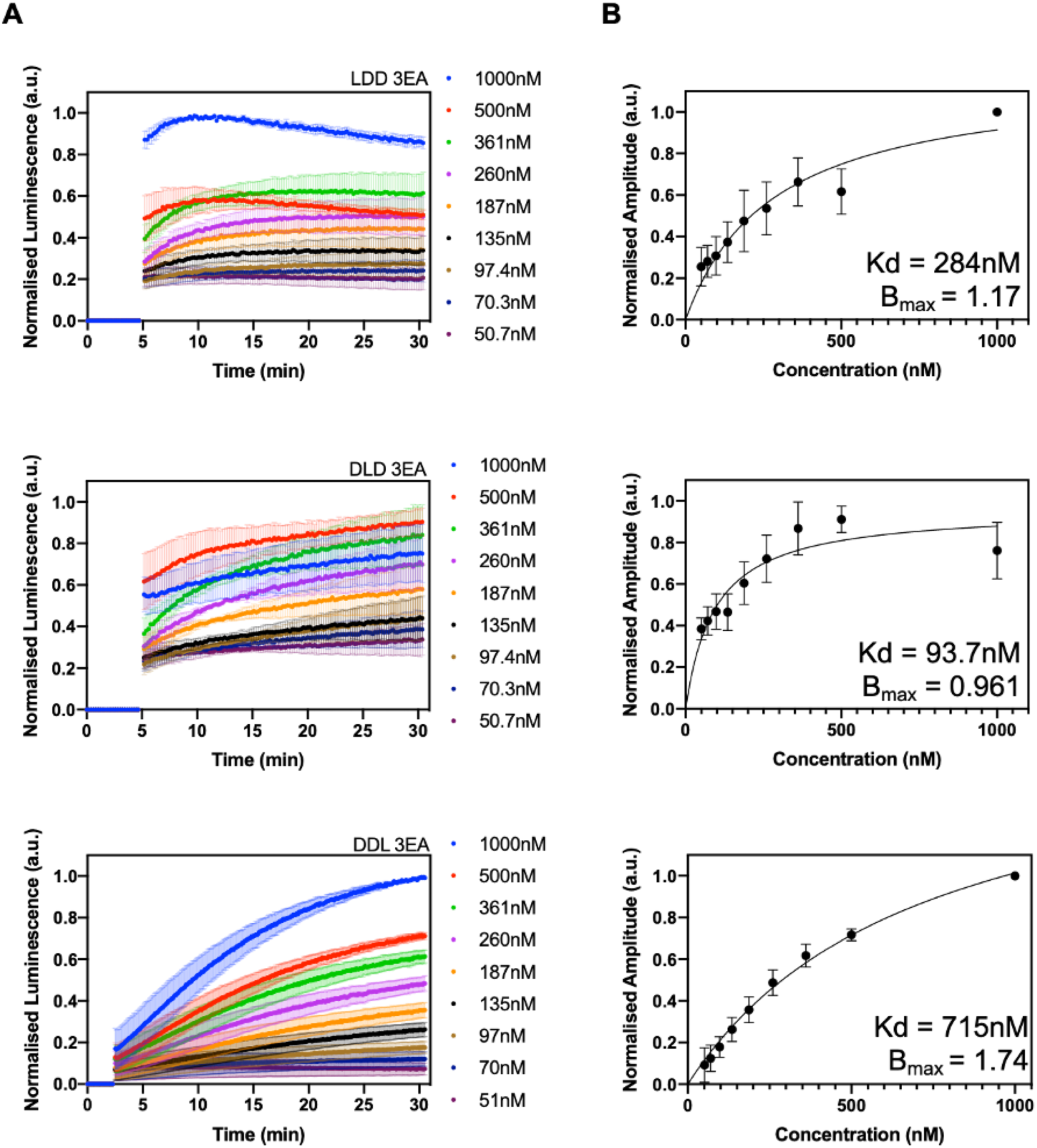
Binding Analysis of the PINK1 3EA Variants to 11S. (**a**) A titration series of the PINK1 LD 3EA precursors was conducted from 51nM to 1000nM and assayed for binding against a fixed concentration of purified 11S, 200pM. (**b**) Normalised maximum amplitudes were calculated and plotted against each concentration of precursor. Data were fit to a simple model for one site binding and the associated K_d_ and B_max_ (maximum amplitude) values were calculated based on the fit. Data represent an N=3 biological repeats and error bars the SEM.

**Figure S13:**
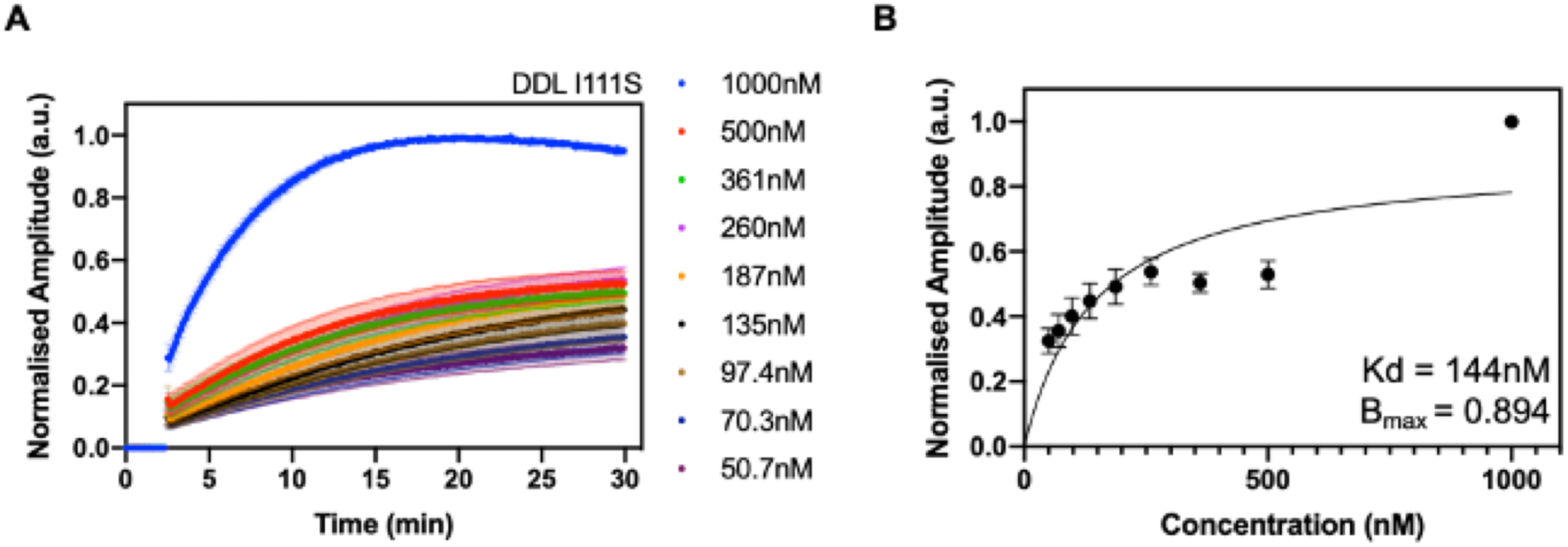
Binding Analysis of the Pathogenic PINK1 Variant I111S to 11S. **(a)** A titration series of DDL I111S precursor was conducted from 50.7nM to 1000nM and assayed for binding against 200pM purified 11S. **(b)** Maximum amplitude at each concentration of I111S was calculated and plotted. Data was fitted to a one-site binding model and associated K_d_ and B_max_ (maximum amplitude) values shown.

**Figure S14:**
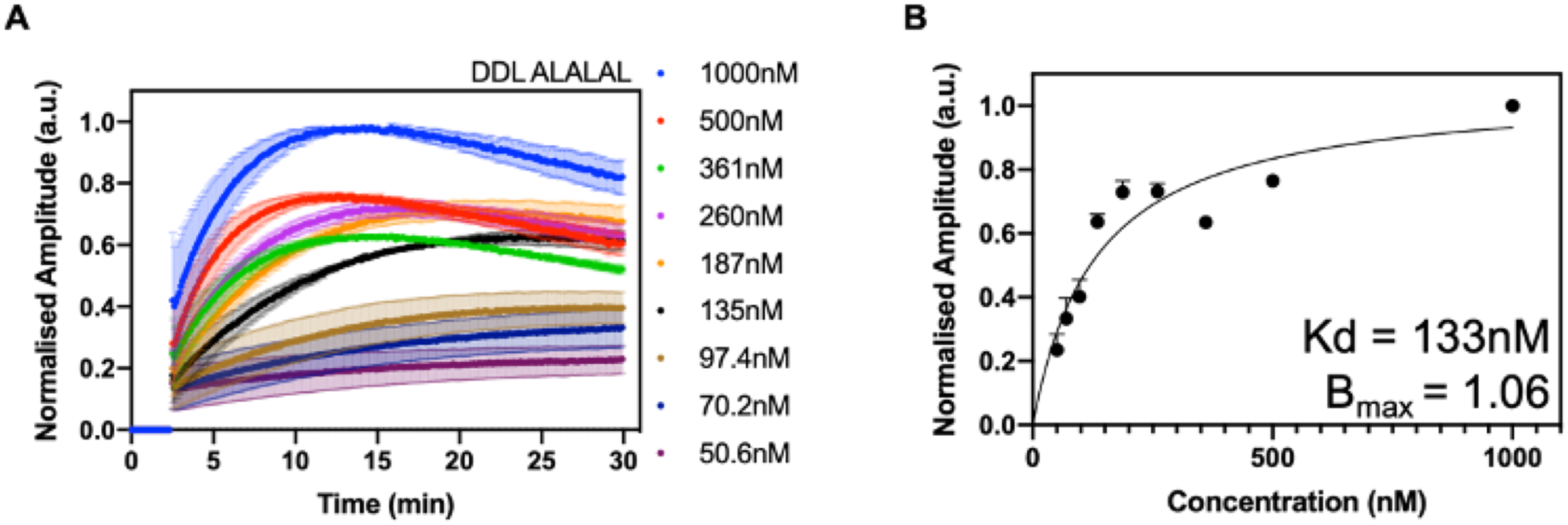
Binding Analysis of the PINK1 ALALAL variant to 11S. **(a)** A titration series of PINK1 ALALAL was conducted from 50.6nM to 1000nM and assayed for binding against 200pM purified 11S. **(b)** Maximal amplitude at each ALALAL concentration was plotted and data fit to a simple one site binding model, resulting K_d_ and B_max_ (maximum amplitude) data are shown.

